# Structural dynamics in the CENP-A nucleosome impacted by protein-protein interactions with centromere protein N^†^

**DOI:** 10.1101/2025.11.10.686994

**Authors:** Abhik Ghosh Moulick, Sylvia Erhardt, Wolfgang Wenzel, Mariana Kozlowska

**Author notes:** Supplementary Information available: [details of any supplementary information available should be included here]. See DOI: 00.0000/00000000.

## Abstract

Noncanonical nucleosomes at the centromere contain the histone variant CENP-A, playing a crucial role in chromosome segregation. CENP-A is highly regulated and its centromeric organization is partially regulated by the centromere protein N (CENP-N). Despite its importance, the protein–protein interactions within the complex formed between CENP-A nucleosomes and CENP-N remain poorly understood at a molecular level. Here, we employ the SIRAH coarse-grained molecular dynamics (MD) simulations to investigate the binding interface and structural rearrangements of proteins in the CENP-A nucleosome complexed with CENP-N. We aim to assess the stability of the CENP-A nucleosome and the change in its plasticity upon the CENP-N binding. By the set of *µ*s-long MDs, we reveal enhanced flexibility in the N-terminal region of CENP-A and stabilization of its RG loop in the complex with CENP-N. This is demonstrated to have rather minor effects on the overall stability of the nucleosome and changing its compactness. Nevertheless, the data suggest the binding of CENP-N allosterically changes the conformational states of CENP-A and impacts its interactions with other proteins in the histone core. A distance-based contact map analysis further elucidates key residues mediating the interaction between CENP-A and CENP-N, while umbrella sampling simulations quantify their binding free energy, which remains challenging to measure experimentally.

## 1 Introduction

In eukaryotic cells, DNA is compacted within the nucleus through a series of structural levels of chromatin, where the nucleosome core particle (NCP) serves as fundamental unit ^1^. NCP typically comprises 147 bp long duplex DNA wrapped approximately 1.7 times around positively charged histone octamer protein complexes, which are spaced by so called linker DNA, that facilitate higher-order structure ^2^. Canonical nucleosomes consist of an octamere of two copies of the four canonical histones H2A, H2B, H3, and H4. Histones are highly conserved and consist of a globular core region with a smaller fraction of disordered, positively charged segments that extend outwards from the core, called the N-terminal histone tails. These tails are highly dynamic, modulated by a large number of post-translational modifications and involved in chromatin organization and chromosome condensation. ^1,3–6^. Further complex processes, including chromatin looping, formation of topologically associating domains, and compartmentalization into chromosome territories, progressively organize the genome, allowing it to be efficiently packed within the nucleus ^7,8^.

Variations in nucleosome composition, differences in DNA sequences, linker DNA length, and various post-translational modifications, influence NCP-NCP interactions, and thereby folding of chromatin, changing its higher-order structure and gene regulation in virtually all biological processes, including DNA repair, replication, and gene expression ^9–11^. For example, cancer-associated mutations in linker histone have been reported to disrupt nucleosome stacking and decompaction of higher-order chromatin structures, which in turn result in irregular gene expression and contribute to oncogenic transformations ^12^. Changes in canonical nucleosomes via histone variants are pivotal in the epigenetic process by shaping the identity of specific regions in the genome, for example, centromeres. Centromeres act as platforms for the assembly of kinetochores ^13,14^, which are large protein complexes that mediate the attachment of spindle microtubules to chromosomes during mitosis and meiosis, maintaining proper chromosome segregation. Centromeric DNA in human cells are composed of repetitive AT-rich *α*-satellite DNA sequences ^15,16^, along with the presence of the histone H3 variant CENP-A. At centromeric chromatin, CENP-A replaces canonical H3 in a subset of nucleosomes, thus defining the site of kinetochore formation. Thereby, CENP-A acts as an epigenetic marker ^17–19^ for centromere localization and kinetochore formation.

In total, sixteen inner kinetochore proteins are associated with centromeric nucleosome, which collectively known as the constitutive centromere-associated network (CCAN) ^20–22^. The specific binding of NCPs containing CENP-A with CCAN occurs by complex protein-protein interactions (PPIs), where CENP-N and CENP-C directly recognize the CENP-A nucleosome necessary for further kinetochore assembly and kinetochore segregation ^23–25^. While CENP-N acts as a reader and locator of CENP-A, anchoring the CCAN complex ^26^ specifically to CENP-A–containing nucleosomes, CENP-C binds CENP-A nucleosomes for recruiting and organization of other CCAN components ^27,28^. Moreover, CENP-N was reported to facilitate the stacking of CENP-A–containing nucleosomes and the formation of nucleosomal arrays through contacts between its *α*6 helix and the DNA of neighboring nucleosomes. This leads to the formation of densely packed chromatin at centromeric regions ^29^. Cryo-EM studies and biophysical analyses further confirmed that these protein–protein and protein–DNA interactions are key elements underlying the formation of higher-order centromeric structures ^29^. Hydrogen/deuterium exchange (HX) coupled to mass spectrometry experiments additionally revealed that the N-terminal domain of CENP-N adopts a folded conformation, with the first 200 residues forming the major interface with CENP-A nucleosomes ^30^. Consistently, structural studies showed that human CENP-N (residue GLU3, THR4, and THR7) confers binding specificity through interactions with the L1 loop of CENP-A, particularly its exposed RG motif (ARG80–GLY81), which are further stabilized by electrostatic contacts with nucleosomal DNA. Several positively charged residues of CENP-N (ARG44, LYS45, ARG11, LYS81, LYS148, and ARG169, ARG170), located proximal to the DNA backbone, likely form stabilizing interactions with the DNA phosphate groups ^25^. Overall, the functionality of such noncanonical nucleosomes depends on their intrinsic structural flexibility ^26,30^ and on conformational changes stimulated by CCAN protein binding and DNA interactions ^25,30–32^. Despite these understandings, detailed dynamical studies of how specific CENP-A–CENP-N interactions guide centromere function at the level of a single nucleosome and beyond remain elusive. Above all, quantitative data on binding energetics are still lacking from both experiments and simulations, yet such information is important for understanding whether CENP-N binding is essentially constitutive or dynamic, and how it compares in strength to other nucleosome-protein interactions. These energetic insights, along with structural observations, help to better understand how protein–protein and protein–DNA interactions regulate nucleosome function at the centromere.

The structual changes of canonical nucleosomes, for example their loop formation ^33,34^, DNA breathing and unwrapping ^6,34–36^, twist defects ^36^, nucleosome sliding ^37^, as well as some structural characteristics of histone variant nucleosomes ^38,39^ have been studied to date through different computational methods ^6,40^ with diverse structural resolutions ^41–43^. Here, molecular dynamics and the use of enhanced sampling simulations ^44,45^ are the most applicable in the field. MD simulations have already been employed for histone variant nucleosome. For example, Kohestani et al. ^38^ elucidated the molecular mechanism by which H2A.B leads to a less compact nucleosome state, thereby increasing genetic accessibility and gene transcription. Bowerman et al. ^46^ reported altered dynamics and allosteric pathways mediated by changes in L1-loop interactions between the two H2A.Z histone copies. Kono et al. ^44^ performed free energy calculation based on unwrapping of superhelical turn of CENP-A-containing nucleosome and revealed that the lowest free energy corresponds to the state where 16 to 22 base pairs were unwrapped. Still, studies related to the interaction of histone variant nucleosomes with other biological macromolecules present in the nucleus remain obscure.

Due to the large system size of NCPs and biological complexes they may form by interacting with other proteins, gaining long-timescale conformational insights and PPIs through atomistic simulations are computationally expensive. Therefore, coarsegrained (CG) models ^47–51^ serve as a useful alternative for exploring such systems. While polymer-based CG models ^50,52^ and mesoscopic models ^43,56^ are often used for nucleosome and chromatin modeling, higher resolution CG force fields (FF) like SIRAH (Southamerican Initiative for a Rapid and Accurate Hamiltonian) ^59,60^ or MARTINI ^61^ are capable to decipher proteinprotein and protein-DNA interactions with a higher accuracy. MARTINI uses largely a bottom-up strategy for bonded interactions and a top-down strategy for non-bonded interactions during parameterization, while SIRAH is mostly bottom-up, physics-based CG FF with long-range electrostatics, thus enabling capturing of hydrogen bond-like interactions and other interactions that are typically absent in less fine FFs. In addition, owing to its parameterization scheme, SIRAH permits unbiased simulation of the secondary structure that often possesses constraints in other models, and it was demonstrated to correctly reproduce PPI ^62^ and protein-DNA interactions ^63–65^ in comparison to experimental observations. Furthermore, the SIRAH FF was recently applied to understand protein-DNA interactions and dynamics of canonical nucleosomes in comparison to all-atom simulations ^65,66^.

Due to adequate quality of SIRAH in describing interactions between biological macromolecules and structural dynamics of biologically relevant complexes ^67–70^, we selected this FF for indepth characterization of protein binding and dynamics of the histone variant nucleosome system with CENP-A towards understanding of its CENP-N–driven structural dynamics and binding energetics. Using the SIRAH FF, we first assess nucleosome stability in the presence and absence of CENP-N and characterize the binding interface between CENP-N, CENP-A, and DNA. We then analyze the conformational flexibility of CENP-N in both its free and bound states, providing microscopic details of its structuring upon NCP binding. Finally, we quantify the PPIs binding strength through umbrella sampling (US) simulations. With this approach, we aim to gain mechanistic insights into how CENP-N contributes to CENP-A nucleosome stability and recognition, providing a better understanding of the functionality and structural changes of centromeric NCPs.

## 2 Methods

### 2.1 System preparation

We considered the cryo-EM structure of CENP-A nucleosome in complex with the CENP-N protein taken from protein data bank (PDB) under ID 7U46^29^. The histone part of this cryo-EM structure has missing histone tails, and CENP-N protein has some missing residues, see Table S1 in Supplementary Information (SI). These missing residues and histone tails were modeled in the present work using AlphaFold3^71^. Further structure refinements were performed by means of atomistic molecular dynamics simulations using Amber99SB-ILDN force field ^72^ and TIP3P water model ^73^. Simulations were performed in the GROMACS simulation package ^74^, version 2019.2. At first, the NCP system was immersed in a cubic box of water of dimensions 167.35*×*167.35*×*167.35 Å^3^. The system was neutralized by adding 149 Na^+^ ions and minimized for 50000 steps using the steepest descent algorithms. Further equilibration proceeded in two steps using NVT, and later NPT, ensembles keeping position restraint on heavy atoms. It was carried out at 300 K and 1 bar using the V-rescale thermostat and Parrinello-Rahman barostat with isotropic pressure coupling. Short-range van der Waals and Coulomb interactions were truncated at 10 Å. Long-range electrostatics were treated using the Particle Mesh Ewald (PME) method with a cubic interpolation order of 4 and a Fourier grid spacing of 1.6 Å. Neighbor lists were updated every 10 steps using the Verlet cutoff scheme with a grid-based search. The production run of all-atom simulation was performed for 10 ns with 2 fs integration time step employing periodic boundary conditions in all directions. All bonds involving H atoms were constrained using the LINCS algorithm. The final structure obtained from the atomistic simulation (presented in Fig.1a) was used for further CG simulations.

### 2.2 Coarse-grained MD simulations

CG-MD simulations were performed for two main NCP systems: (i) CENP-A nucleosome with bounded CENP-N protein (this system was labeled as “CENP-A Nucleosome + CENP-N”) and (ii) CENP-A nucleosome without CENP-N protein (labeled as “CENP-A Nucleosome”). The second system was prepared by removing CENP-N from the final structure obtained from the atomistic simulation of CENP-A Nucleosome + CENP-N (see Section 2.1). Simulations were performed following the computational setup in the previous work of simulating canonical NCP system ^66^. All-atom refined structures were protonated using PDB2PQR ^75^ server at neutral pH assumption following the AMBER naming scheme necessary for SIRAH. The structures obtained were mapped into CG form using SIRAH 2.2 tools (Fig. 1b). After mapping, the complexes were solvated in a PBC box with SIRAH WT4 water type ^76^. All systems were neutralized and the physiological conditions were modeled by adding Na^+^ and Cl^-^ ions at a salt concentration of 0.15 M. The required number of ions, PBC box size, and number of solvent CG beads are listed in Table 1. The simulation box was defined using a truncated octahedral shape with a 20 Å distance between the solute and the box edges to ensure that the complexes did not interact with their periodic images. Energy minimization was performed in two steps: (i) first, for 50000 steps with the protein side chains minimization while restraining the backbone, and (ii) second, for 5000 steps with the entire system minimized without constraints. In both cases, the steepest descent algorithm was utilized. The side chain equilibration enhances structural stability of protein molecule by avoiding significant distortions to the secondary structure. Subsequently, the solvent molecules were equilibrated around the complex by running a 5 ns simulation with harmonic restraints applied to all CG beads. The temperature of the system was maintained at 300 K using a V-rescale thermostat ^77^ and the constant isotropic pressure of 1 bar using Parrinello-Rahman barostat ^78^, respectively. Before production run, an additional run of 10 ns was performed keeping restrain on protein, allowing the DNA to relax in the interface. Finally, unrestrained production run was carried out for 10 *µ*s. The time step for the simulation was maintained at 20 fs. The electrostatic interactions were computed using the PME method with a 12 Å cutoff and a grid spacing of 2 Å. Van der Waals interactions were treated with a 12 Å cutoff. Three separate replicas of each system were simulated for a duration of 10 *µ*s by starting from different random seeds. The solvated CENP-N protein without the NCP was simulated for three independent replicas using the same protocol as for other complexes, excluding the DNA relaxation step. The system specific information is specified in Table 1.

**Table 1.** Summary of the systems setup used for the CG-MD simulations with SIRAH.

| System studied | CENP-A Nucleosome + CENP-N | CENP-A Nucleosome | Only CENP-N |
| --- | --- | --- | --- |
| PBC Box (Å) | 192.1×181.3×157 | 171.3×161.5×139.8 | 115.4×108.8×94.2 |
| No. of CG beads | 67454 | 48078 | 14788 |
| No. of CG solvent molecules | 14620 | 10222 | 3269 |
| No. of Na<sup>+</sup> ions | 536 | 399 | 101 |
| No. of Cl<sup>-</sup> ions | 387 | 247 | 104 |
| Salt concentration (M) | 0.15 | 0.15 | 0.15 |

**Fig. 1.**
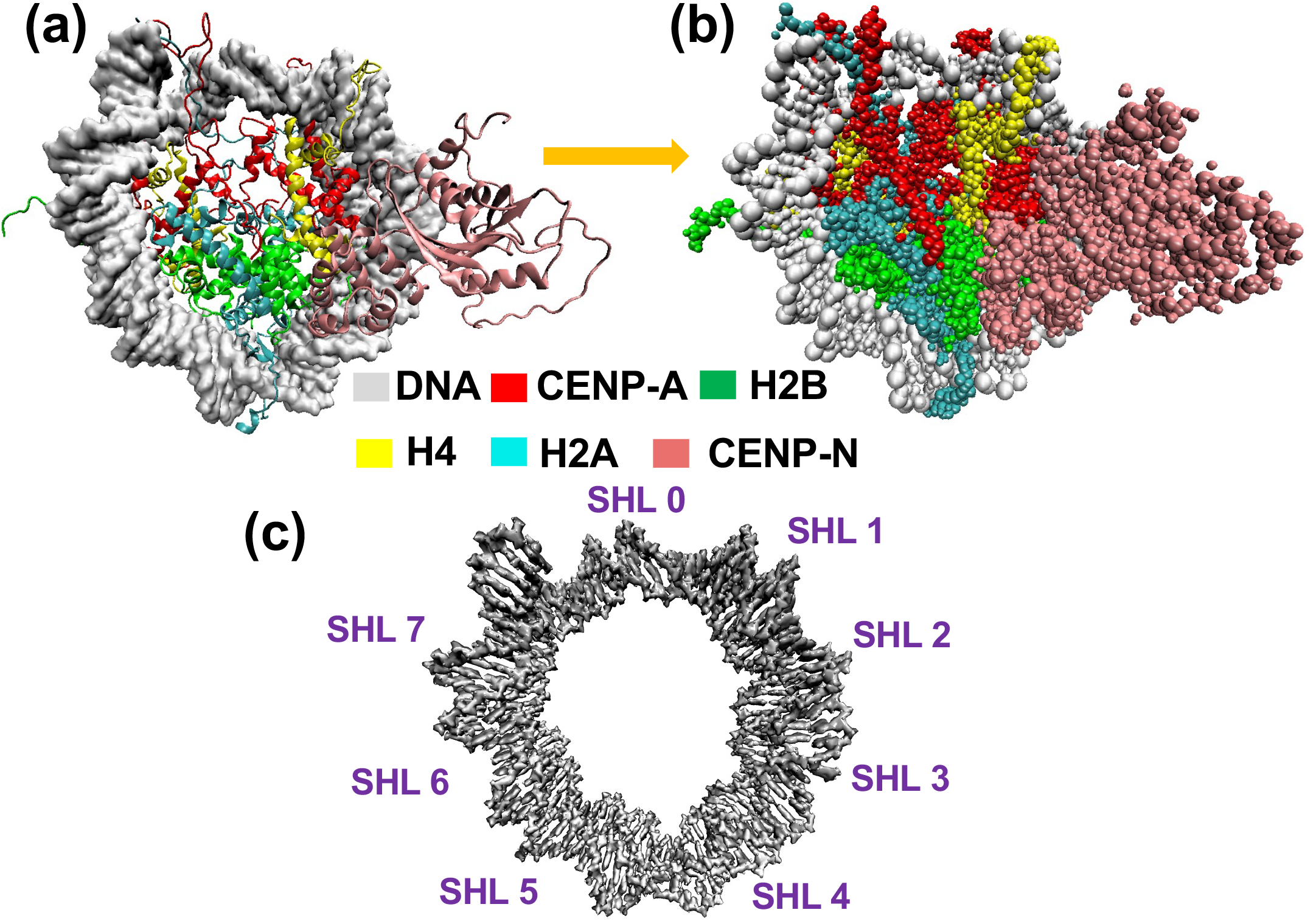
(a) The all-atom structure of the CENP-A-containing nucleosome along with the CENP-N protein obtained in cryo-EM ^29^ and further refined using atomistic simulations (this work). The histone core and CENP-N protein are visualized using New Cartoon and four histone pairs are marked in red, green, yellow and cyan for CENP-A, H2B, H4 and H2A, respectively, while DNA is visualized using QuickSurf and is marked in grey. The CENP-N protein is marked in salmon. (b) The coarse-grained representation of the CENP-A nucleosome with bonded CENP-N as mapped from its refined all-atom structure. (c) The visualization of superhelical location (SHL) sites marked over nucleosomal DNA. Each SHL typically spans approximately 10 base pairs, ranging from SHL0 to SHL ±7.

SIRAH-Backmap tools ^79^ were used for backmapping CG-MD trajectories into atomistic resolution. The backmapping procedure involves reconstructing atomistic positions on a per-residue basis ^80^, preserving the geometric structure (internal coordinates), followed by protonation and minimization using the atomistic force field ff14SB atomistic force field ^81^ in Amber-Tools ^82^ tleap module.

### 2.3 Analysis of nucleosome dynamics

To characterize the dynamical properties of the CENP-A nucleosome and its structural changes upon the specific protein binding simulated with CG-MD, we applied a set of analyses explained here. Global stability was monitored through the root-mean-square deviation (RMSD), while local flexibility was captured by root-mean-square fluctuations (RMSF). For both analyses, we considered the backbone CG beads of both histone and DNA. RMSD and RMSF were calculated relative to the energy-minimized structure explained in Section 2.2. Changes in the overall compaction of the nucleosome were quantified using the radius of gyration (Rg), which is a mass-weighted root-mean-square distance of all beads from the center-of-mass (COM). To probe sequence- and region-specific histone–DNA interactions, we computed contact maps between protein and DNA beads. Following earlier work ^83^, native contacts were defined when the specified beads (GC for protein representing carbon atom, PX for DNA representing backbone phosphate atom) were within 7 Å. In the SIRAH CG representation, the direct identification of specific interactions such as hydrogen bonds or salt bridges is not feasible due to the reduced resolution of the model. Instead, residue–residue interactions were characterized using this distance-based contact definition. For each residue pair, the average contact population value was calculated as the fraction of simulation frames in which the contact was present. An average contact value of 1.0 indicates that the residues remained in contact throughout the entire trajectory, representing a highly stable interaction. Contacts with values below 0.4 were classified as transient or weak, indicative of higher flexibility in that region. The average contact values between protein residues were further visualized in the form of 2D contact maps, where the axes correspond to residue indices and the color scale indicates the stability of the contacts across the trajectory. A dark/intense color (the average contact value close to 1.0) shows that the two residues stayed in contact throughout the trajectory (stable, persistent interaction) while lighter color (less than 0.4) shows weak or transient contacts. All analyses were performed using GRO-MACS tools, while MDAnalysis was utilized to calculate the contact map ^84^.

The secondary structure of proteins was calculated using SIRAH secondary structure tool. It uses the positions of CG backbone beads to approximate the backbone geometry defined as a structured element in terms of helical and extended regions (i.e. *β* -strands), or unstructured element defined as a coil. Since the SIRAH CG geometry of proteins retains enough information about local backbone curvature and spacing, these CG-based geometric rules provide an accurate distinction between helices from sheets ^59^. To extract dominant modes of motion in histone CENP-A upon CENP-N binding and to construct free energy landscapes (FEL), representing thermodynamic map of conformational space of a protein, the principal component analysis (PCA) was employed. It is capable to reveal the functionally relevant conformational states of molecules during the trajectories and reduce dimensionality that helps in identifying configurational spaces with a limited number of degrees of freedom. In this method, a 3*N ×* 3*N* covariance matrix of positional fluctuations of CG beads relative to every other coordinates over time is constructed based on simulated trajectory. Thus, it captures the correlated motions of coordinates, and the diagonalization of this matrix yields eigenvectors, representing principal motion directions, and the corresponding eigenvalues, indicating the magnitude of fluctuations along these directions. The trajectory is then projected onto these eigenvectors to derive the principal components (PCs). The first two PCs describe the dominant large-scale motions and are oftenemployed to construct a two-dimensional FEL. It is based on the estimation of the joint probability density function (P(PC1,PC2)) obtained from a histogram of PCs and defined as *ΔG*_*FEL*_(*x, y*) = −*k*_*B*_*Tln*(*P*(*x, y*)*/P*_*max*_), where *P*_*max*_ is the probability of the most probable state, *k*_*B*_ is Boltzmann constant and T is the temperature. The PCA and FEL were calculated using MDAnalysis Python library ^84,85^ providing input coordinates of CG beads. Together, the analyses performed provide a comprehensive view of nucleosome stability, flexibility, as well as collective dynamics.

The errors for the quantities calculated were estimated where applicable. The error of the average is given by 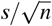, where *s* is the standard deviation of the mean values over the replicas, and *n* represents the number of replicas.

### 2.4 Umbrella Sampling simulations

To understand PPIs and binding free energy between CENP-A protein in an NCP and CENP-N CCAN protein, US simulations using SIRAH FF were performed ^62,86^. The initial structure of the complex was taken from the final structure obtained from the atomistic simulations as discussed above. It was placed in a box of 320 *×* 200 *×* 160 Å^3^, so that the complex was aligned parallel to the X-axis. The system was solvated, neutralized, and minimized as per the prior simulation protocol, explained in Section 2.2. Equilibration was then performed under NVT and NPT conditions, maintaining consistency with the MD simulation protocol. To initiate the US simulation, at first we performed steered molecular dynamics (SMD) simulation to generate initial conformations for US windows. The CENP-N protein was pulled away from its initial position near the CENP-A side of the NCP into the bulk solvent along the X-axis, applying a harmonic pulling potential with a force constant of 1000 kJmol^*™*1^ nm^*™*2^ with a pulling rate of 0.0001 nm/ps. To prevent drifting of the system along the reaction coordinate, a harmonic restraint with force constant of 20 kJmol^*™*1^ nm^*™*2^ for whole nucleosome, except CENP-N protein, was applied. Positions of the molecular system simulated were saved during the course of pulling with 60 structures generated as US windows for separate MD simulations. Each window structure was saved every 0.1 nm from the initial position of CENP-N and up to a COM distance between the CENP-A and CENP-N proteins of 7.95 nm. From a COM distance of 8.05 to 14.65 nm, the spacing was fixed at 0.2 nm. The schematic of US procedures is depicted in Fig. S1.

Each system in the US window was independently further simulated for 25 ns in thr NPT ensemble followed by an additional MD simulation for 50 ns with a V-rescale thermostat and Parrinello–Rahman barostat. A harmonic bias potential with a force cosntant of 1000 kJmol^*™*1^ nm^*™*2^ was applied to each window. This value was selected within the range (500–2000 kJ/mol *·* nm) reported in previous SIRAH umbrella sampling studies ^62,63^ on protein–protein interactions and was further optimized to achieve adequate histogram overlap between adjacent windows, ensuring sufficient sampling across the reaction coordinate. Fig. S2 depicts overlap of histograms from different windows obtained along reaction coordinate. Finally, the potential of mean force (PMF) was derived using the weighted histogram analysis method (WHAM) ^87,88^ as implemented in GROMACS to eliminate the influence of the applied bias. Statistical errors were estimated by using default Bayesian bootstrapping algorithm built into the WHAM program ^88^.

The binding free energy (*ΔG*) between proteins, indicating the strength and the character of PPIs was calculated as the difference in free energy between the bound and unbound states. The unbound state was defined as a conformational state where interactions between the proteins are completely diminished, corresponding to a plateau (near-zero interaction) in the PMF plot. *ΔG* can be calculated mathematically as follows:

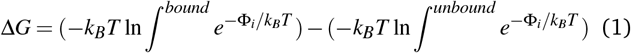

Φ_*i*_ represents the PMF value associated with the *i*^th^ bin along the reaction coordinate. The error in the binding energy is calculated by propagating the errors ^89^ from the minimum PMF (bound state) and the plateau region (unbound state) using the following equation:

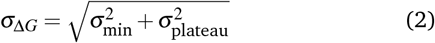

Here *σ*_*min*_ is the error at the minimum PMF and *σ*_*plateau*_ is the average error in the plateau region.

## 3 Results

### 3.1 Stability of CENP-A nucleosome upon CENP-N binding

Since the function of NCPs depends on their structural flexibility, interactions with other NCPs or molecules, as well as upon changing environment conditions, we aimed to analyze the stability of CENP-A nucleosomes in the presence and absence of CENP-N pro-tein. For that, we calculated RMSD and Rg using data obtained for three independent replicas (see more details in Fig. S3 and Fig. S4). The RMSD has been calculated for both the histone protein core and nucleosomal DNA separately. We further categorized the RMSD calculations for the histone core based on the inclusion and omission of the residues present in histone tails, which are generally inherently flexible, thus, hindering the understanding of subtle changes in other protein regions. The respective data are labeled as “with tails” and “without tails” in Fig. 2. The time evolution of RMSD for all cases is presented in Fig. S3 while time dependent RMSD of only histone tails are depicted in Fig. S5 .

**Fig. 2.**
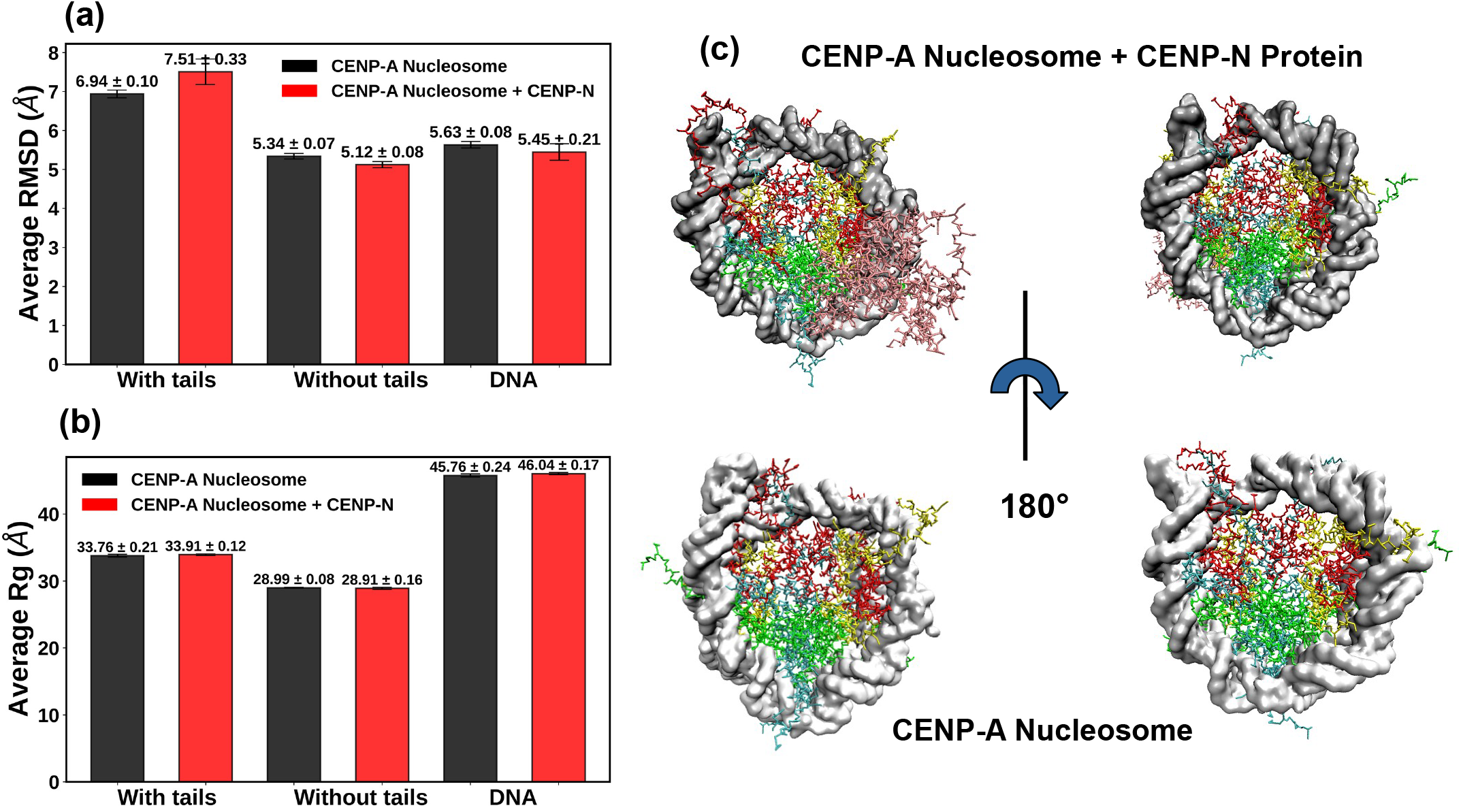
(a) The average root mean square deviation and (b) radius of gyration of the nucleosome with and without histone tails considered and the nucleosomal DNA alone. Structural parameters for the CENP-A nucleosome with and without CENP-N protein are marked in red and black, respectively. (c) The structure of the systems simulated after 10 *µ*s CG-MD simulations: the CENP-A nucleosome + CENP-N protein complex (top), and CENP-A nucleosome (bottom). Two views of the same complex are depicted.

The average RMSD plot (Fig. 2a) shows that considering histone tails in the RMSD calculation, the fluctuation is higher in the system with bounded CENP-N to the CENP-A system (RMSD of 7.51*±*0.33 Å, see marked in red) as compared to the unbound CENP-A system (6.94*±*0.10 Å, in black). It is connected to the interactions induced by the presence of CENP-N explained further. The RMSD of systems without the consideration of the histone tails is lower, indicating higher histone core stability. In addition, the RMSD of the histone core in the presence of CENP-N is even slightly smaller, i.e., 5.12*±*0.08 Å while in the absence of CENP-N the average RMSD is 5.34*±*0.08 Å. This indicates a possible stabilization of the nucleosome upon CENP-N binding. The average value of RMSD of nucleosomal DNA in the presence and absence of CENP-N is 5.45*±*0.21 Å and 5.63*±*0.08 Å, respectively (Fig. 2a). It suggests a slight rigification of some base pairs that are located near the CENP-N binding site, permitting their smaller structural deviations. Together with a slight histone core structural stabilization, it provides evidence for possible NCP plasticity changes, which we discuss in the following section.

The average radius of gyration for both systems is depicted in Fig. 2b. The presence of CENP-N protein does not alter larger structural moves and the shape of the NCP. Time evolution of Rg is shown in Fig. S4. The final structure of the NCPs after the simulation is visualized in Fig. 2c. The upper panel depicts the CENP-A nucleosome with bound CENP-N, while the lower panel shows the unbounded CENP-A NCP. Distinct orientations are displayed for each system to convey the complete molecular architecture and assembly. The figure demonstrates that the systems considered in this study retained their structural integrity throughout the microsecond-scale CG-MD simulations.

### 3.2 CENP-N binding interface: contacts with CENP-A and DNA

CENP-N interacts specifically with the CENP-A histone and nucleosomal DNA, forming extensive contacts that are critical for its binding specificity and centromere function. Fig. 3a highlights the interaction interface between DNA, CENP-A, and CENP-N. The L1 loop of CENP-A is shown in red, while the DNA region, spanning SHL 2 to 3.5 (see Fig. 1c) that participates in binding, is highlighted in yellow. Residues of CENP-N interacting with CENP-A and DNA are shown in green and blue, respectively. This representation provides a structural overview of the binding sites identified from experimental cryo-electron microscopy analysis ^25^. To evaluate the stability of experimentally observed residue contacts under dynamic conditions, we analyzed these interactions over 10 *µ*s of CG-MD simulations, from which 8 *µ*s were considered for data analysis.

**Fig. 3.**
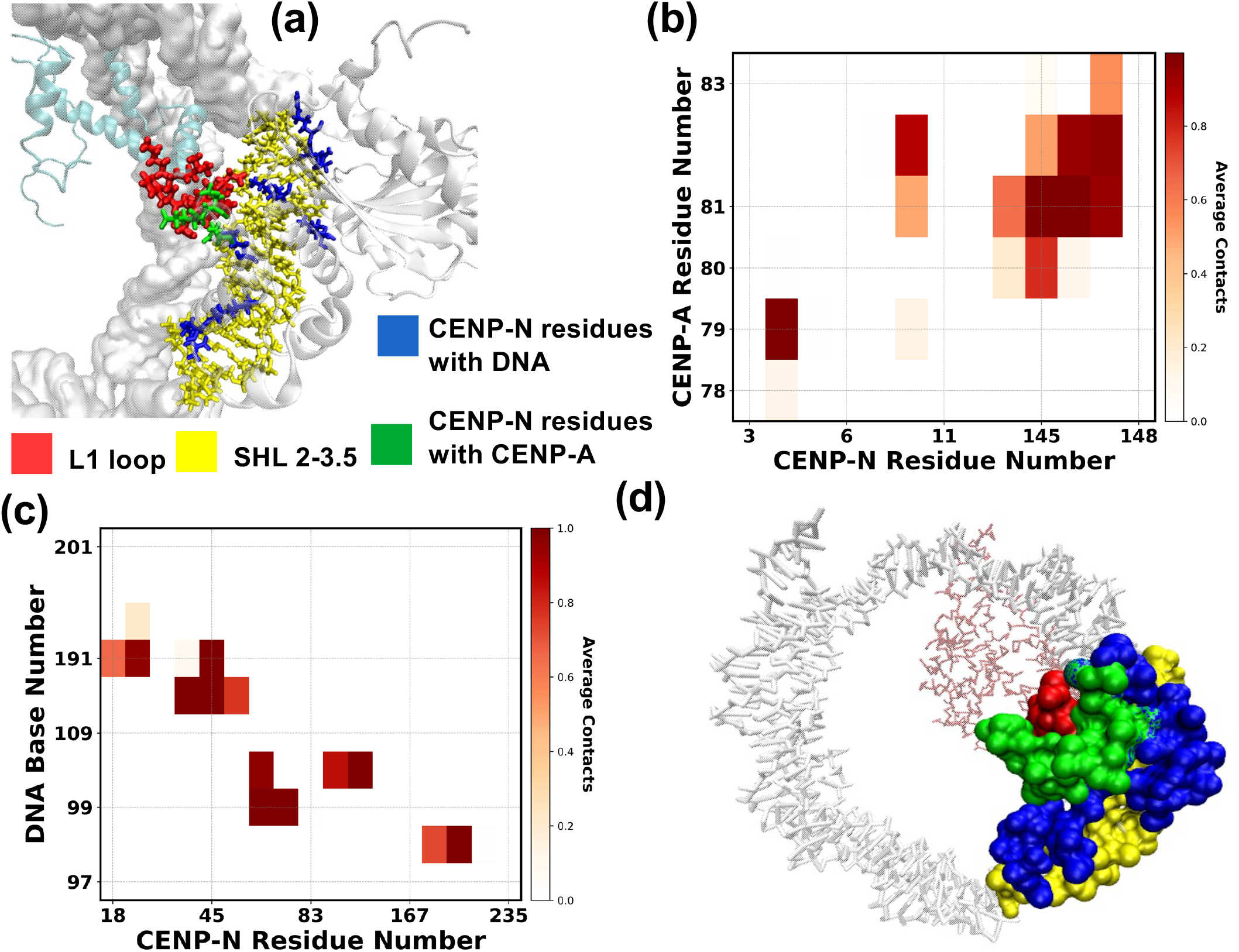
(a) The binding interface involving residues from CENP-A, CENP-N, and DNA bases was obtained from the cryo-EM structure. Red indicates the L1 loop of CENP-A; yellow highlights DNA from SHL 2 to 3.5, the region interacting with CENP-N. Green denotes the CENP-N amino acids that interact with CENP-A, while blue marks those interacting with the DNA SHL. (b) Contact map showing interactions between CENP-N and CENP-A residues. (c) Contact map showing interactions between CENP-N residues and DNA bases. (d) Binding interface obtained from CG simulation with color coding as in panel (a).

In Fig. 3b, the contact map between CENP-N and CENP-A residues over 4000 snapshots from CG-MD simulations is depicted. This contact analysis (see Methods section for details) confirms that CENP-A residues forming the RG loop (i.e., ARG80–GLY81) interact with CENP-N, consistent with cryo-EM data, and suggests stable contacts at this interface during CG-MD. ARG80 of CENP-A interacts strongly with residue ASN145 of CENP-N, resulting in the average contact value of 0.79. Its inter-action with residues PRO144 and GLN146 from CENP-N is weaker (contact value of 0.21 and 0.11 respectively), indicating a broader contact region. Residue GLY81 stands out with strong and multiple contacts, notably with residues ASN145 and GLN146 (both with an average contact population of 1.00), and significant interactions with residues PHE8, PRO144, and TYR147, pointing to a highly engaged interface. The average contact population values of all participating residue pairs are listed in Table S2.

To maintain its main function, CENP-N should primarily bind to the CENP-A protein in the histone core ^23,90^. However, it additionally shows the interaction with the DNA as depicted in Fig. 3a, therefore, the respective contact map between the CENP-N protein and the DNA was calculated (see Fig. 3c). The CG-MD simulations denote several persistent contacts between the DNA bases and CENP-N residues. DNA base G98 exhibits strong interactions with ARG169 and LEU168 of CENP-N, with average contact population of 0.99 and 0.73, respectively. The high frequency of contact formation indicates the attractive character and stability of the CENP-N binding at this interface (see Table S3). Similarly, G99 exhibits stable contacts with LYS81 and VAL82, both showing maximum contact values of 1.00, along with a weaker interaction with TRP83. The adjacent base A100 also forms strong contacts, particularly with LYS148 (1.00), TYR147 (0.85), and again with LYS81 (0.97), indicating a consistent role of this region in DNA recognition. A43 base from DNA strand-2 (DNA base number 190 in Fig.3c) strongly interacts with ARG44, LYS45, and GLU46, with contact values exceeding 0.75, while C44 (DNA base number 191 in Fig.3c) forms multiple contacts with MET18, ASN19, and LYS45, further supporting weak interaction with ARG44. T49 ((DNA base number 196 in Fig.3c)) also shows a modest contact with ASN19. These interactions (see values listed in Table S3) highlight key regions of CENP-N that stably associate with the DNA, suggesting their role in mediating nucleosome binding and stabilizing the DNA–protein interface in the CENP-A nucleosome. The binding interface formed by CENP-A, CENP-N, and DNA is depicted in Fig. 3d, which represents the final snapshot taken after 10 *µ*s-long MD simulation. The amino acid residues from the proteins as well as the DNA nucleic bases are shown in surface representation, using the same color scheme as described in Fig. 3a.

### 3.3 CENP-A structural stability upon CENP-N binding

The CG-MD simulations indicate that the overall structure of the CENP-A-containing NCP remains largely unchanged upon CENP-N binding (Fig. 2). To examine potential structural changes specifically in the CENP-A protein within the histone core in the presence and absence of CENP-N, the RMSF of CENP-A was calculated (see Fig. 4a). The region near the L1 loop of CENP-A (specifically residues CYS75, VAL76, LYS77, PHE78, THR79, ARG80, GLY81, VAL82) possesses lower fluctuations in the CENP-N bound state (data in red) compared to the unbound state (data in black), indicating that CENP-N binding stabilizes this region. Fluctuations in the C-terminal residues (Res 134-Res 140) remain largely unchanged. However, the N-terminal residues exhibit higher value of RMSF in the presence of CENP-N (exceeding 1 Å). Such an observation supports that CENP-N binding induces increased flexibility at the N-terminal region, potentially facilitating conformational adjustments required for further stability. We marked the CENP-A residues showing higher and lower RMSF in the presence of CENP-N relative to the CENP-N unbound state in cyan and green, respectively. These residues are shown in Fig. 4b (with respective regions indicated in Fig. 4a), which represents the final snapshot taken after 10 *µ*s-long MD simulation.

**Fig. 4.**
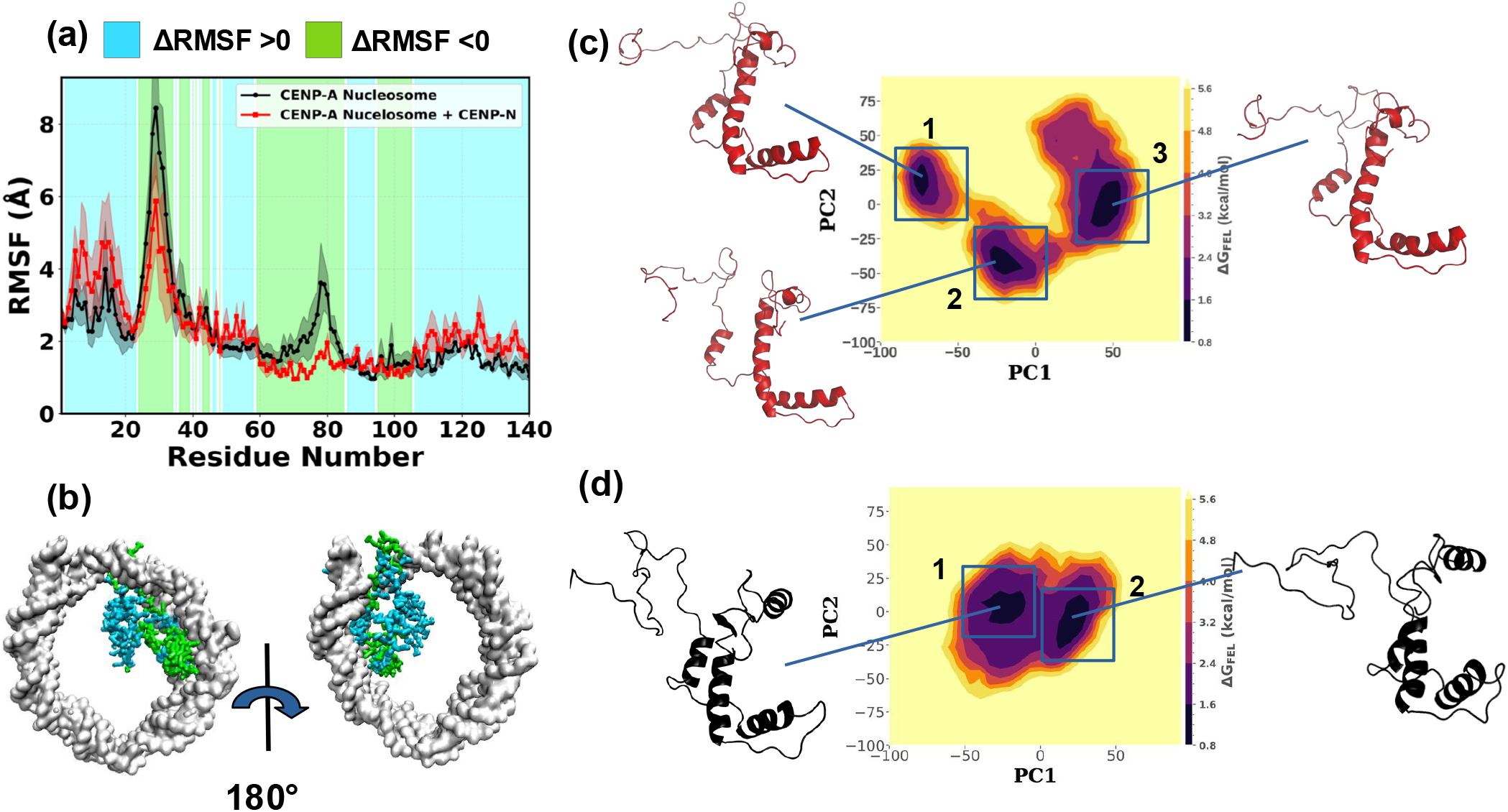
Root mean square fluctuations of the CENP-A protein in the presence (red) and absence (black) of the CENP-N protein. RMSF values were averaged over three independent replicas for both systems. The cyan-shaded areas denote CENP-A residues that exhibit higher RMSF in the presence of CENP-N, while the green-shaded areas mark residues with lower RMSF. These residues are further highlighted in (b), where the final CG-MD structure is depicted with the color coding consistent with the RMSF plot. The free energy landscape of CENP-A along PC1 and PC2 shown in the presence (c) and absence (d) of CENP-N. Representative structures of CENP-A corresponding to each minimum are depicted in panels outside the figure. *ΔG*_*FEL*_ is provided in kcal/mol.

To further understand the CENP-A conformational change upon the CENP-N binding, we performed the PCA analysis as described in the Methods section. In Fig. 4c,d, the FEL plots along two principal components (PC1 and PC2) for CENP-A histone in the presence as well as absence of CENP-N are visualized. In both cases, we observe distinct energy basins indicating distinct stable conformational states of CENP-A. In the absense of CENP-N, the two stable conformational states of CENP-A are structurally similar and are separated by the low energy barrier. However, the number of basins upon the CENP-N binding increases (Fig. 4c), representing stronger structural changes. Such conformational changes likely reflect the specific interactions between CENP-A and CENP-N, stabilizing additional conformational states and revealing a more complex free energy landscape. The identified representative structures corresponding to each energy basin for both cases are depicted in panels outside in Fig. 4c,d. They were identified by picking conformations at the minima of the FEL spanned by PC1 and PC2 shown in Fig. 4c,d. We used backmapped allatom structures to better visualize the conformational changes across the different basins. We see that the stable helical regions remain largely unchanged between structures, while the loop regions exhibit distinct conformations in each minimum. In the CENP-N unbound case (Fig. 4d), the representative structures of CENP-A also show variations in loop conformations while maintaining the helical structure.

### 3.4 Free and bound states of CENP-N

To explore whether CENP-N and CENP-A coevolve to support NCP’s function in the centromere region, we analyzed the structural dynamics of CENP-N both in its complex with the CENP-A-containing nucleosome (labeled as bound CENP-N) and in its isolated state (labeled as free CENP-N). Hence, we performed an additional CG-MD simulation of free CENP-N as explained in the Methods section. Fig. 5 shows the set of analyses conducted to demonstrate the structural differences. Using RMSF calculation depicted in Fig. 5a, differences in fluctuations of specific regions of CENP-N in its both states are visualized. The RMSF plot clearly demonstrates that residues 1–200 become more ordered (stabilized) upon nucleosome binding (see data in red), whereas residues 201–295 display consistently higher RMSF values in both states. Thus, in the bound case, CENP-N exhibits increased flexibility in its C-terminal residues res216-res255 (cyan colored region in Fig. 5a) which are known to be disordered according to previous experimental studies ^30^. In contrast, the N-terminal residues display lower RMSF values in the bounded state of CENP-N, indicating significant stabilization due to interactions with CENP-A and nucleosomal DNA. The N- and C-terminal residues are highlighted in the final conformation of CENP-N in both the bound and free states in panels outside Fig. 5a. Both the CG representation, as well as the backmapped all-atom structures of the protein are visualized. In the CG representation, the CG beads of each amino acid are colored according to their RMSF values as indicated in the color bar. Regions of the molecule with relatively higher RMSF values, indicating greater flexibility, are shown in red, while regions representing relatively lower fluctuations are shown in blue. The RMSF coloring of CENP-N was generated using a user-defined color scale in VMD ^91^. This coloring reflects relative differences in flexibility across the protein rather than absolute RMSF values. The N- and C-terminal residues are marked within dashes circles in the CG representation, while they are highlighted in pink over the protein’s secondary structure in the atomistic representation. The N-terminal residues 1–50 of CENP-N show a more intense red color of the CG beads in the free state of the protein, indicating their higher fluctuations. In the bound state, these fluctuations are significantly reduced, showing medium relative RMSF. In contrast, the C-terminal residues (201–295) display similar color patterns in both free and bound states, suggesting comparable fluctuations, which result in a higher flexibility of this region present in both states of CENP-N.

**Fig. 5.**
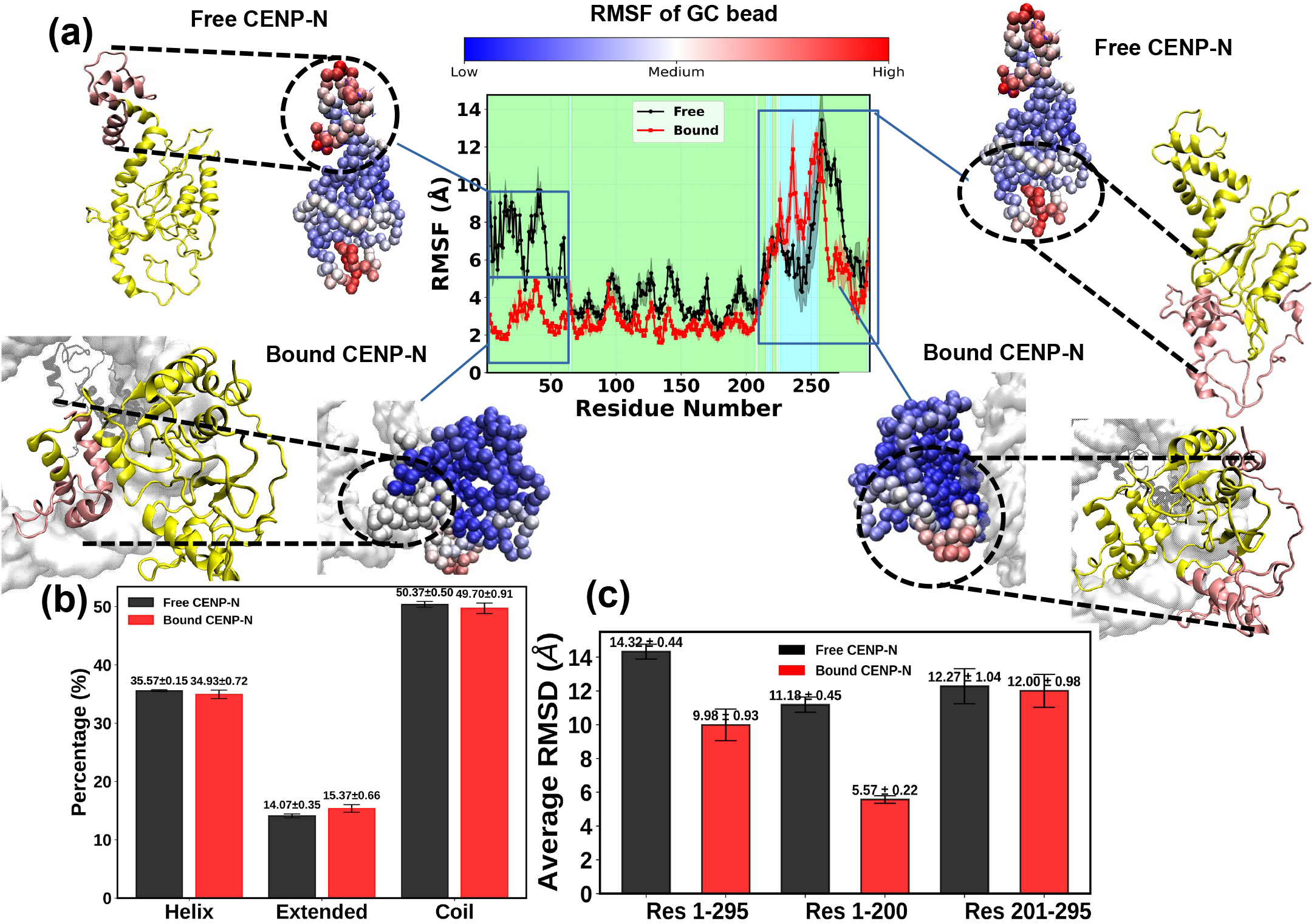
(a) Root mean square fluctuations of the bound (red) and free (black) CENP-N protein, averaged over three independent replicas. . The cyan-shaded areas denote CENP-N residues that exhibit higher RMSF in bound state, while the green-shaded areas mark residues with lower RMSF with respect to free state. Final structures of both free and bound CENP-N are shown at CG and backmapped all-atom resolutions. In all-atom representation, the N- or C-terminal residues are highlighted in pink, while the rest of the protein is shown in yellow. b) Percentage of secondary structure elements observed over the simulation trajectory for both bound and free CENP-N. (c) Average RMSD plots of unbound and free CENP-N for three regions: overall, residues 1–200, and residues 201–295. Error bars represent standard deviations from three independent replicas.

The backmapped all-atom structures reveal that several residues from the N-terminus of CENP-N adopt loop-like conformations in both states, despite largely belonging to helical regions. By contrast, the C-terminal tail predominantly forms loop structures in both bound and free states. Henceforth, it is important to calculate the secondary structure (ss) elements of the CENP-N protein in both the bound and free states across MD simulation replicas. At first, we performed the ss analysis for protein at both free and bound state using SIRAH ss tool as explained in Methods section. Fig. 5b shows the average ss content, along with error bars from three independent replicas, for both systems. We also calculated the time evolution of the ss content for the bound and free states of CENP-N (see Fig. S6) over the last 8 *µ*s. Both results revealed no significant changes in the ss between the two states. Considering the differing flexibility of particular regions of the protein, we identified three different regions that were studied separately in more detail. The first region was located between residue 1 to 50, which showed the strongest changes in the RMSF behavior upon binding. The second region (residue 51 to 200), demonstrating no significant RMSF changes upon binding and the third region (residue 201 to 295), which remain disordered indicating higher RMSF in both bound and free state (Fig. 5a). The secondary structure for these three different regions is given in Table 2, where helix and extended regions represent stable ss, while coil represents unstructured ss. From these data, we see that the first region shows increased contribution of stable ss from 54.07% to 57.70% (especially for helix content, which changes from 49.37% to 52.67%) upon binding as compared to the free state of the protein. In the other regions, the stable ss content versus coil does not change markedly: less than 1% of stable ss decreases or remains unchanged, with overall flexibility largely unaffected upon binding with CENP-A and nucleosomal DNA.

**Table 2.** Secondary structure content (in %) for CENP-N protein regions in bound and free states. Errors are indicated as mean *±* error.

| Category | Res 1-50 |  | Res 51-200 |  | Res 201-295 |  |
| --- | --- | --- | --- | --- | --- | --- |
|  | Bound | Free | Bound | Free | Bound | Free |
| Stable ss: Helix | 52.67±3.75 | 49.37±2.37 | 30.67±1.01 | 32.13±1.67 | 30.60±0.99 | 33.07±2.20 |
| Stable ss: Extended | 5.03±0.88 | 4.70±2.36 | 23.07±1.90 | 22.30±1.22 | 8.20±1.56 | 5.73±0.95 |
| Stable ss: Helix + Extended | 57.70±3.85 | 54.07±3.45 | 53.74±2.16 | 54.43±2.06 | 38.80±1.84 | 38.80±2.40 |
| Coil | 42.30±3.96 | 45.97±1.60 | 46.23±1.31 | 45.57±1.19 | 61.17±0.67 | 61.17±1.80 |

Since experimental studies, such as HX exchange ^30^, have reported that the N-terminal domain of CENP-N adopts a folded conformation upon interaction with CENP-A, particularly involving the first 200 residues, we examined the combined dynamics of the ordered region (residues 1–200) and the disordered region (residues 201–295) in CENP-N through RMSD calculation. At first, we calculated the RMSD of whole CENP-N in its free and bound states, see Fig. 5c. The overall RMSD was significantly higher for free CENP-N than compared to the bound CENP-N with the average RMSD of 14.32*±*0.44 Å and 9.98*±*0.93Å respectively. At the same time, ordered residues (res 1-200) show a decrease in RMSD from 11.18*±*0.45Å to 5.57*±*0.22 Å upon nucleosome binding, whereas the flexibility of the disordered region remains similar in both conditions. To further assess the compactness of CENP-N considering residues in these different regions, we calculated the corresponding Rg (see Fig. S7). Rg is slightly smaller in the bound state of CENP-N, however, the average Rg for all three cases (entire CENP-N, the ordered region, and the disordered region), indicates less differences than the average RMSD. Thus, while region-specific RMSD analysis reveals local stabilization of the N-terminal residues upon nucleosome binding, the overall Rg remains largely similar, indicating that CENP-N retains its global size and shape while undergoing local conformational adjustments.

### 3.5 CENP-N binding free energy

To quantify the strength of the PPIs between the CENP-A protein in the NCP and the modulating CENP-N protein, the binding energy was calculated using the US method. Here, we considered the reaction coordinate, *ξ*, as the COM distance between the CENP-A protein and the COM of the ordered part of CENP-N (res 1-200). The choice of the ordered region of CENP-N was guided by the RMSF and RMSD analyses depicted in Fig. 5a,c, which showed that residues 201–295 remain disordered irrespective of nucleosome binding. Focusing on the ordered regions ensures that the sampled conformational space reflects meaningful interactions and avoids contributions from highly flexible segments. The resulting PMF plot along with the calculated error bars is presented in Fig. 6a. The binding energy of the complex can be calculated from the difference between the highest and lowest values of the average PMF curve (details in the Method section). Thus, the binding free energy between CENP-A and CENP-N is concluded to be -7.92±0.99 kcal/mol.

**Fig. 6.**
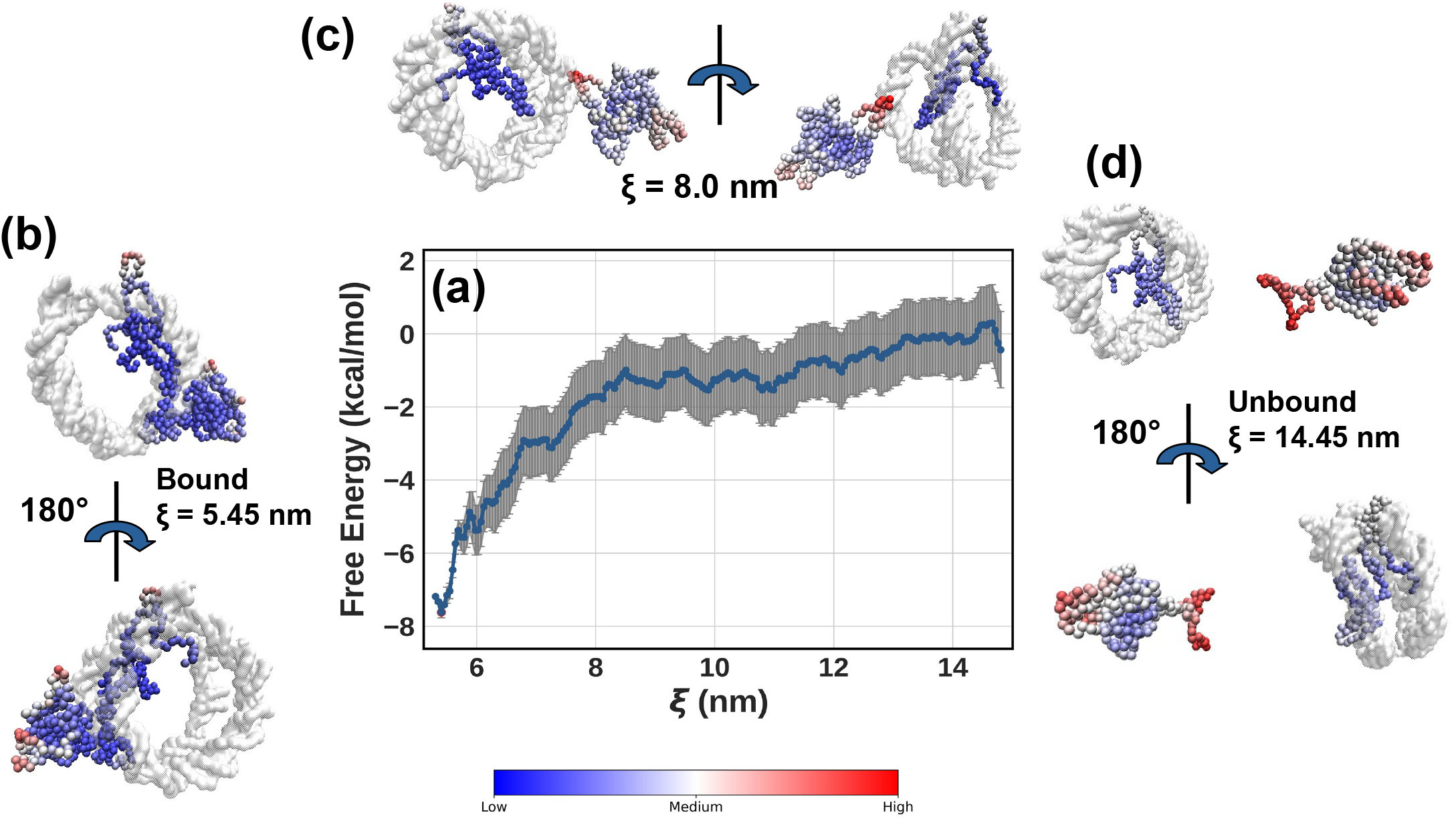
(a) The PMF curve representing the binding free energy between the CENP-A and CENP-N proteins in the centromere NCP obtained using the umbrella sampling method. The error bars calculated using Bayesian bootstrapping algorithm are depicted in grey. The 3D representation of the binding site and the proteins involved is depicted using CG beads, along with the nucleosomal DNA depicted in white surface representation. Different structures representing the COM reaction coordinate, *ξ*, of: (b) *ξ* = 5.45 nm, (c) *ξ* = 8.0 nm, and (d) *ξ* = 14.45 nm, are visualized for clarity. Bead colors represent the relative RMSF fluctuations, where red beads indicate regions of CENP-A and CENP-N with the highest fluctuations, while blue beads indicate the smallest fluctuations. The RMSF coloring was generated using a user-defined color scale in VMD ^91^. Two views of the same system are depicted at the selected COM reaction coordinate.

To visualize the structural changes of CENP-A and CENP-N at three different *ξ* values, the representative structures of the system calculated were further inspected. To do so, the CG beads of all residues in both proteins were visualized and colored according to their relative RMSF values, calculated from the restrained trajectories at the corresponding *ξ* value. A more intense red color indicates significant fluctuations of molecular entity representing the bead, while a more intense blue color represents smaller fluctuations. At *ξ* = 5.45 nm, corresponding to the bound state between the proteins, we observe higher fluctuations in the disordered region of CENP-N that is not involved in the direct protein-protein contact (Fig. 6b). At *ξ* = 8.0 nm, fluctuations at both terminals of CENP-N increase (Fig. 6c), while in the unbound state (*ξ* = 14.45 nm) they display much stronger fluctuations (Fig. 6d). The unbound state of CENP-N essentially represents the free state of CENP-N, where a substantial portion of the protein exhibits a disordered nature. For CENP-A, fluctuations increase in the unbound state compared to the bound state, although the increase is less significant compared to CENP-N since the protein is bounded inside the histone core. The higher structural stability of both proteins upon their binding is clearly visible from this analysis and complements the dependencies observed in the unbiased MD simulations.

## 4 Discussion

The plasticity of the centromeric NCP is a key factor in facilitating its function ^92^. Its interaction with binding proteins directs chromatin folding and the compaction of genetic information, processes that are essential for accurate chromosome segregation and the maintenance of genome stability ^11^. Here, we examine the binding of CENP-N and the resulting structural changes in the NCP. Overall, our study investigates the PPIs within the NCP, with a particular focus on how CENP-N binding influences the dynamics and stability of the CENP-A nucleosome. Because nucleosome dynamics occur over extended timescales, we employed a CG-MD to capture these interactions and evaluate how the underlying protein–protein associations contribute to nucleosome functionality.

The RMSD analysis of NCPs without histone tails reveals no significant structural differences upon the CENP-N binding with rather small structural stabilization of 0.22 Å (Fig. 2a), indicating that the core histone fold remains structurally stable irrespective of the CENP-N presence. However, when histone tails are included in this analysis, a modest increase in RMSD in the presence of CENP-N is observed, suggesting that the histone tails may adopt more dynamic or altered conformations upon CENP-N binding. Interestingly, the radius of gyration remains largely unchanged across all comparisons, including those performed for histone cores with and without tails and the DNA (Fig. 2b). This indicates that despite local flexibility changes, particularly in the histone tails, the overall compactness of the nucleosome remains constant. These results suggest that the role of the CENP-N protein is not to further compact individual nucleosomes, but rather to facilitate inter-nucleosomal interactions that were reported to drive centromeric chromatin folding and organization ^29^.

Starting with the binding interface involving DNA bases, CENP-A amino acids and CENP-N amino acids reported in cryo-EM ^29^, we performed several replica MD simulations each of 10 *µ*s, which revealed stable residue-residue contacts between the NCP and CENP-N. PPIs between CENP-A and CENP-N were shown to demonstrate a high specificity of the interaction interface enriched in polar, charged, and aromatic side chains, suggesting diverse binding interactions. The RG loop of CENP-A within its L1 loop, ARG80 and GLY81 emerge as central points of contact in the simulation based on the average contact values (see Tab. S2), which were also suggested by the cryo-EM study of Chittori et al. ^25^. Considering the chemical nature of the individual contributing residues, we found that ARG80 of CENP-A, with its positively charged guanidinium group, interacts strongly with ASN145 of CENP-N and moderately with PRO144 and GLN146. These residues possess polar or partially polar side chains, enabling electrostatic and hydrogen bond-based interactions. Even if the SIRAH CG FF cannot properly treat subtle hydrogen bonds, it is known to adequately represent electrostatic interactions, which are essential in proper mimicking the PPIs of the complex studied. Notably, GLY81 of CENP-A exhibits a high degree of flexibility and forms extensive contacts with ASN145, GLN146, PHE8, PRO144, and TYR147 of CENP-N, supporting the idea that this position serves as a hub for both polar and hydrophobic interactions. THR79, a polar uncharged residue of CENP-A, interacts strongly with THR4 of CENP-N, with an average contact value of 0.99 (see Table S2), likely forming a hydrogen-bond network via their side-chain hydroxyl groups. Additionally, a hydrophobic VAL82 participates in both polar (ASN145, GLN146) and aromatic (PHE8, TYR147) contacts with CENP-N, suggesting van der Waals and hydrophobic interactions may also contribute. Moreover, a negatively charged ASP83 forms notable contacts with polar aromatic TYR147 (average contact value 0.55), which are likely mediated through hydrogen bonding or electrostatic attraction. These findings indicate that the binding interface between CENP-A and CENP-N is stabilized through a combination of diverse interactions, which together contribute to a high PPIs specificity and strength of the CENP-A/CENP-N interaction, which was calculated to be -7.92±0.99 kcal/mol based on umbrella sampling simulations performed in this work. Interestingly, while cryo-EM studies ^25^ have previously identified GLU3, THR7 as participating in the interaction interface between CENP-N and the RG loop of CENP-A, the contact-based analysis performed based on CG-MD simulations in the present work did not reveal stable or significant interactions involving these residues.

The interaction between the nucleosomal DNA and CENP-N was also probed using a distance cutoff approach. The DNA bases, primarily belonging to SHLs 2 to 3.5, and amino acids from CENP-N involved in the interaction are listed in Table S3. This observation is consistent with previous experimental studies ^25^. The simulation suggests that several arginine and lysine residues from CENP-N, such as ARG169, LYS81, LYS148, ARG44, and LYS45, interact with DNA, indicating that the interaction is predominantly electrostatic. The distance-based interaction analysis also identified additional amino acids likely to participate in DNA binding. For example, LEU168 and VAL82 were detected to interact with DNA through hydrophobic contacts, while GLU46 and TYR147 frequently contact DNA bases (average contact value 0.76 and 0.85 respectively), suggesting potential hydrogen-bond-like interactions in the coarse-grained representation. MET18 and ASN19 are also recognized as potential DNA binding residues due to their ability to form hydrophobic interactions and hydrogen bonds, respectively.

To monitor any allosteric changes in the histone core upon binding of CENP-N, we focused on the conformational arrangement of the CENP-A protein in the presence and absence of CENP-N. The RMSF analysis of CENP-A in Fig. 4a revealed sequence-dependent differences in the structural flexibility upon CENP-N binding. The L1 loop region of CENP-A exhibits reduced RMSF in the presence of CENP-N, whereas in the absence of CENP-N, this region displays markedly increased flexibility. This suggests that binding of CENP-N confers structural stabilization to the L1 loop, potentially through direct interactions. RMSF analysis further shows that the N-terminal region of CENP-A exhibits higher flexibility when bound to CENP-N compared to its unbound state. This suggests that interaction with CENP-N enhances the dynamic behavior of the CENP-A N-terminus. In parallel, contact map base analysis (Fig. S8) revealed a reduction in the average number of contacts formed by residues 13–22 of CENP-A with histone H2A when CENP-N is present (Fig. S8a). Conversely, in the absence of CENP-N, this region engages in more persistent contacts with H2A (Fig. S8c) with participating residues marked within a circle. Overall, this study indicates that the binding of CENP-N allosterically disrupts contacts between the CENP-A N-terminal tail and histone H2A, thereby enhancing the conformational freedom of the N-terminus.

The enhanced fluctuations of the CENP-A N-terminal tail (see Fig. 4a) observed in our simulations suggest that this region relies on interactions with specific binding partners for protein stabilization when bound to CENP-N. This dynamic behavior is consistent with experimental observations ^93^, highlighting the functional importance of the N-terminus of CENP-A for centromere functionality and kinetochore assembly. For example, the N-terminus of CENP-A was shown to contribute to the stabilization of the centromere binding protein CENP-B by direct interaction ^94,95^. Moreover, it acts as a recruiter of key kinetochore proteins such as CENP-C and CENP-T at both ectopic sites and endogenous centromeres in Schizosaccharomyces pombe and human cells ^96,97^. Thus, the enhanced fluctuations of the N-terminal tails of CENP-A, as captured by RMSF in Fig. 4a, may serve as a signature of its active engagement in modulating interactions essential for centromere function.

Furthermore, CENP-A’s unstructured N-terminal tail bears post-translational modifications ^98^. Several studies have demonstrated that for instance phosphorylation of serine 7, 16 or 18 within the CENP-A N-terminal domain significantly influences centromeric chromatin structure and function ^99–101^. These residues lie near the region that shows altered contact behavior in CG simulations (see residue 13 to 22 in Fig. S8), suggesting that CENP-N binding may modulate the exposure or accessibility of phosphorylation sites, thus indirectly impacting downstream chromatin remodeling or signaling events (see Fig. S8 a). Therefore, the observed destabilization of local contacts (Fig. S8 a) and increased flexibility of the N-terminus of CENP-A (residue 13 to 22 in Fig. 4a) upon CENP-N binding is not only structurally plausible but also biologically meaningful, potentially contributing to regulating centromere function via a modulation of post-translational modification, recruitment of kinetochore proteins, or higher-order chromatin structure formation.

To understand the molecular determinants of centromere assembly, it is essential to quantify the binding energetics of the CENP-A–CENP-N complex. Binding energy not only reflects overall stability but also reveals how different regions of a multivalent protein contribute to interaction, providing mechanistic insight that cannot be captured by static structural measurements alone. Although, calculating binding energy experimentally for this nucleosome-like system is challenging to date due to the multivalent and context-dependent nature of their interactions.

Finally, the calculated binding free energy between the CENP-Acontaining NCP and the CENP-N protein from CG-MD simulations in our work captures the strength and complexity of the interaction (Fig. 6a), while further analyses of trajectories generated uncovers the split nature of CENP-N: the N-terminal domain becomes ordered upon binding to the NCP, whereas the C-terminal region remains largely disordered (Fig. 6b-d). A similar observation was also reflected during the independent CG-MD simulation of CENP-N in its bonded and a free state as discussed in Section 3.4. RMSF base study supports the split nature of CENP-N (see Fig. 5a), where N- and C-terminals of CENP-N remain disordered in the free state, while the N-terminal becomes ordered in the bound state with the NCP. Such a localized structural ordering in our simulations aligns well with experimental HX exchange data that analyzed the structural dynamics of the N-terminal domain of CENP-N bound to CENP-A nucleosome, as well as in its free state ^30^. The experimental study reported substantial HX pro-tection throughout the N-terminal with up to 200 residues upon binding to the CENP-A nucleosome, indicating a transition to a more stable and ordered conformation. In our work, MD simulations show that the C-terminal region of CENP-N (Res 201-295) remains largely disordered, consistent with the experimental observation that residues 209–240 are not involved in binding ^25^. Notably, the increased order in the bound state in our simulations does not involve major changes in the secondary structure (Fig. 5b) for the whole protein, with no global alterations observed over multiple replicas. However, regional differences are evident: the N-terminal segment (residues 1–50) exhibits increased stable secondary structure upon binding, suggesting its role in mediating interactions with CENP-A and nucleosomal DNA. The middle segment (residues 51–200) maintains a balanced distribution of structured and unstructured elements (see Table 2, showing minimal response to binding. By contrast, the C-terminal region (residues 201–295) is predominantly disordered in both states, consistent with its reported higher flexibility. Despite the local changes described, the overall Rg of CENP-N remains largely unchanged (a decrease of around 1.07*±*0.86 Å was obtained, see Fig. S7), indicating that the global size and shape of the protein are mostly maintained. Taken together, these findings indicate that binding of CENP-N to the NCP primarily stabilizes the N-terminal region of CENP-N, while the central and C-terminal regions remain largely unaffected. This study suggests that the binding-induced conformational changes in CENP-N are driven by localized ordering rather than large-scale structural rearrangements.

## 5 Conclusions

In this study, we provide molecular insights into the interaction between the CENP-A nucleosome and one of its key binding partners, CENP-N, using a combination of biased and unbiased molecular dynamics simulations. Notably, the binding interface predicted by cryo-EM studies remains conserved in the coarse-grained simulations performed, linking structural and dynamical perspectives of this biological complex. In addition, CENP-N association enhances conformational fluctuations in the N-terminal region of CENP-A, a feature likely to facilitate further protein–protein interactions essential for centromere assembly and function. A split nature of CENP-N was observed in unbiased MD simulations, as well as upon binding to the nucleosome core particle revealed by umbrella sampling method. The multitude of analyses and dependencies reported in this work supports diverse experimental observations, thus demonstrating the SIRAH coarse-grained force field reliably captures the long microsecond-scale dynamics of this centromeric complex.

Finally, binding free energy analyses provided a quantitative perspective on CENP-A–CENP-N association in NCPs, revealing insights into promising directions for the regulation of centromere function through post-translational modifications, chromatin remodelers, or the rational engineering of synthetic centromeres. Together, our findings provide a critical mechanistic understanding in the characterization of protein-protein and protein-DNA interactions, deepening our perception of microscopic processes in centromeres and offering a foundation for future studies aimed at targeted control of centromere functionality. Subsequent studies may further explore experimental measurements of binding energetics to complement and validate the computational predictions presented here.

## Author contributions

Abhik Ghosh Moulick: Writing – review and editing, Writing – original draft, Visualization, Validation, Software, Methodology, Investigation, Formal analysis, Data curation, Conceptualization. Sylvia Erhardt: Writing – review and editing, Validation, Funding acquisition. Wolfgang Wenzel: Writing – review and editing, Resources, Funding acquisition. Mariana Kozlowska: Writing – review and editing, Supervision, Resources, Project administration, Methodology, Funding acquisition, Formal analysis, Conceptualization.

## Data availability

Data generated is available from the corresponding author upon reasonable request.

## Acknowledgements

This research was made possible by funding from the Carl-Zeiss-Stiftung and Center SynGen. The authors gratefully acknowledge the computing time provided on the high-performance computer HoreKa by the National High-Performance Computing Center at KIT (NHR@KIT). This center is jointly supported by the Federal Ministry of Education and Research and the Ministry of Science, Research and the Arts of Baden-Württemberg, as part of the National High-Performance Computing (NHR) joint funding program (https://www.nhr-verein.de/en/our-partners). HoreKa is partly funded by the German Research Foundation (DFG).

