## Supplementary Information for "Structural dynamics in the CENP-A nucleosome impacted by protein-protein interactions with centromere protein N^†^"

Table S1: Missing residues in the histone core and base pairs of the DNA in PDB 7U46 that were modeled in the present work using AlphaFold3 for each chain. Chains J and K correspond to DNA.

| **Chain** | **Modeled Residues** |
|---|---|
| A (Histone CENP-A) | Res 1–45, Res 135–140 |
| B (Histone H4) | Res 1–23, Res 103 |
| C (Histone H2A) | Res 1–14, Res 114–130 |
| D (Histone H2B) | Res 1–33, Res 126 |
| E (Histone CENP-A) | Res 1–44, Res 136–140 |
| F (Histone H4) | Res 1–24, Res 103 |
| G (Histone H2A) | Res 1–15, Res 116–130 |
| H (Histone H2B) | Res 1–33, Res 126 |
| I (CENP-N) | Res 92–98, Res 213–295 |
| J (DNA) | Base 1, Base 147 |
| K (DNA) | Base 1, Base 147 |

Table S2: Structural contacts between the residues of CENP-A and CENP-N calculated from MD trajectories. The average contact value is provided in the parenthesis. An average contact value of 1.0 indicates that the residues remained in contact throughout the entire trajectory, representing a highly stable interaction. Contacts with values below 0.4 were classified as transient or weak, indicative of higher flexibility in that region

| **Residue (CENP-A)** | **Residue (CENP-N)** |
|---|---|
| PHE78 | THR4 (0.12) |
| THR79 | THR4 (0.99), PHE8 (0.15) |
| ARG80 | ASN145 (0.79), PRO144 (0.21), GLN146 (0.11) |
| GLY81 | ASN145 (1.00), GLN146 (1.00), PHE8 (0.49), PRO144 (0.63), TYR147 (0.94) |
| VAL82 | PHE8 (0.88), ASN145 (0.51), GLN146 (0.95), TYR147 (0.95) |
| ASP83 | TYR147 (0.55) |

Table S3: Structural contacts between the DNA bases and CENP-N residues participating in the binding interface formation. Average contact value is in the parenthesis. They were obtained with the same approach as explained in Table S2 and the Methods section in the main body.

| **DNA Base** | **CENP-N Residues** |
|---|---|
| G98 (DNA strand 1) | ARG169 (0.99), LEU168 (0.73) |
| G99 (DNA strand 1) | LYS81 (1.00), VAL82 (1.00), TRP83 (0.01) |
| A100 (DNA strand 1) | TYR147 (0.85), LYS148 (1.00), LYS81 (0.97) |
| A43 (DNA strand 2) | ARG44 (1.00), LYS45 (1.00), GLU46 (0.76) |
| C44 (DNA strand 2) | MET18 (0.65), ASN19 (0.96), LYS45 (1.00), ARG44 (0.09) |
| T45 (DNA strand 2) | ASN19 (0.22) |

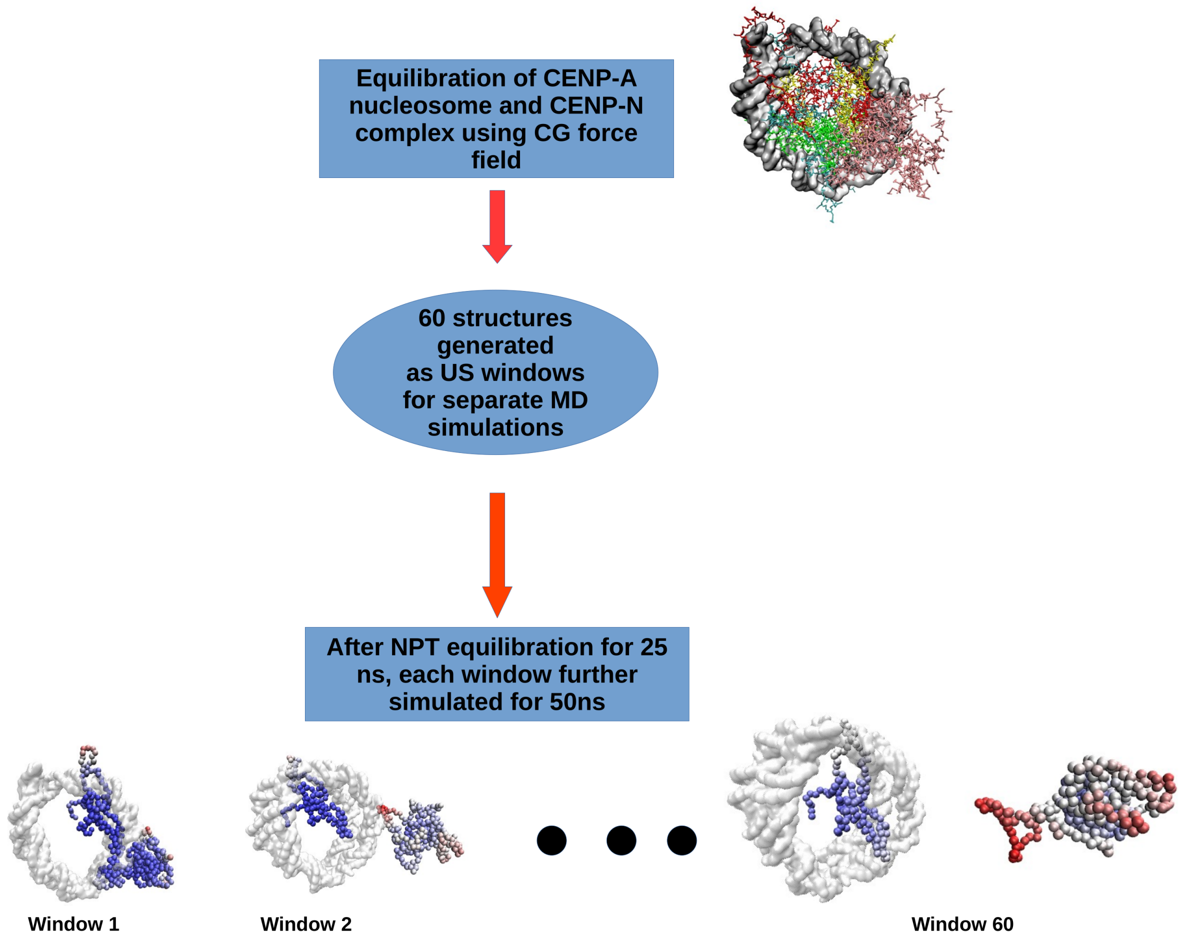


Fig. S1: The schematic workflow applied for umbrella sampling calculations used in this study.

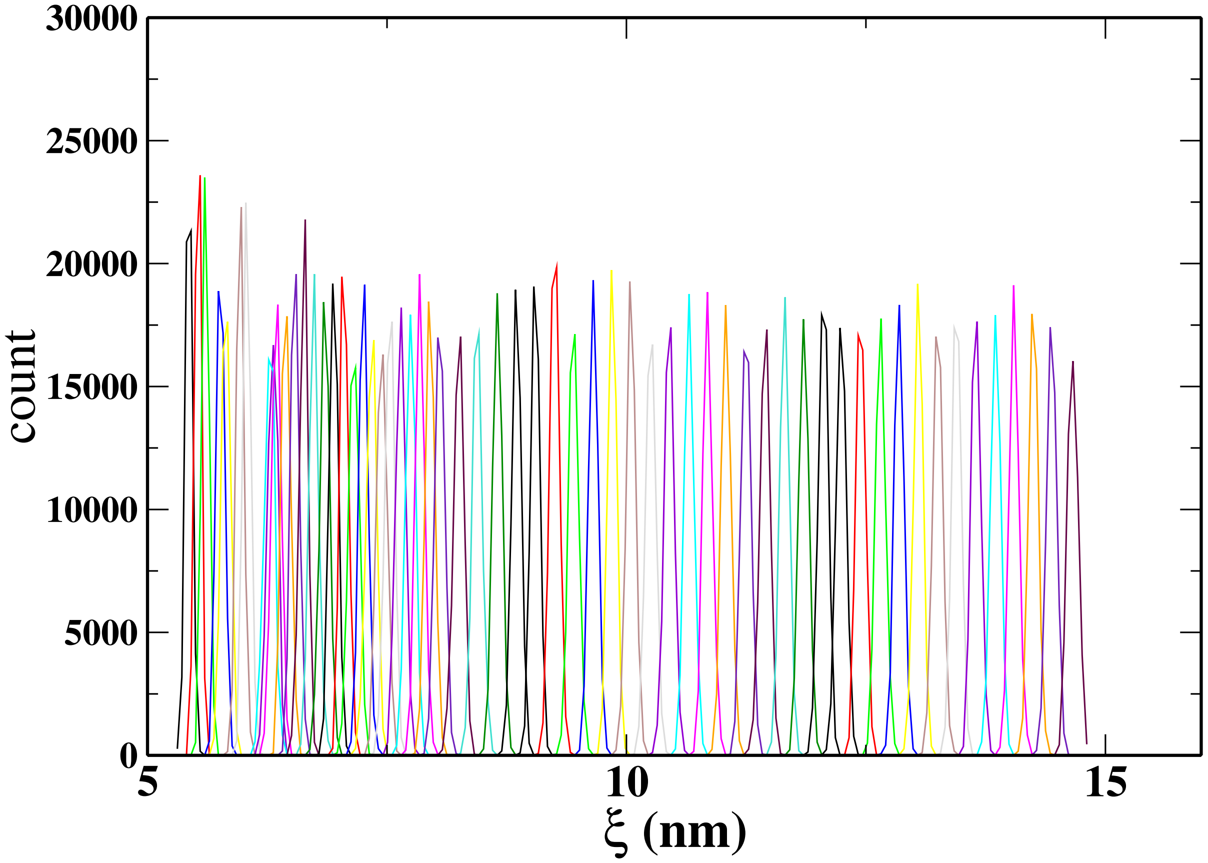


Fig. S2: Overlap of histograms from umbrella sampling (US) windows along the reaction coordinate $\xi$, defined as the COM distance between CENP-A and the ordered region (residues 1–200) of CENP-N. US windows were generated from steered MD, where CENP-N was pulled away from the CENP-A side of the NCP along the X-axis using a harmonic potential. Sixty windows were spaced every 0.1 nm up to 7.95nm and every 0.2 nm up to 14.65 nm.

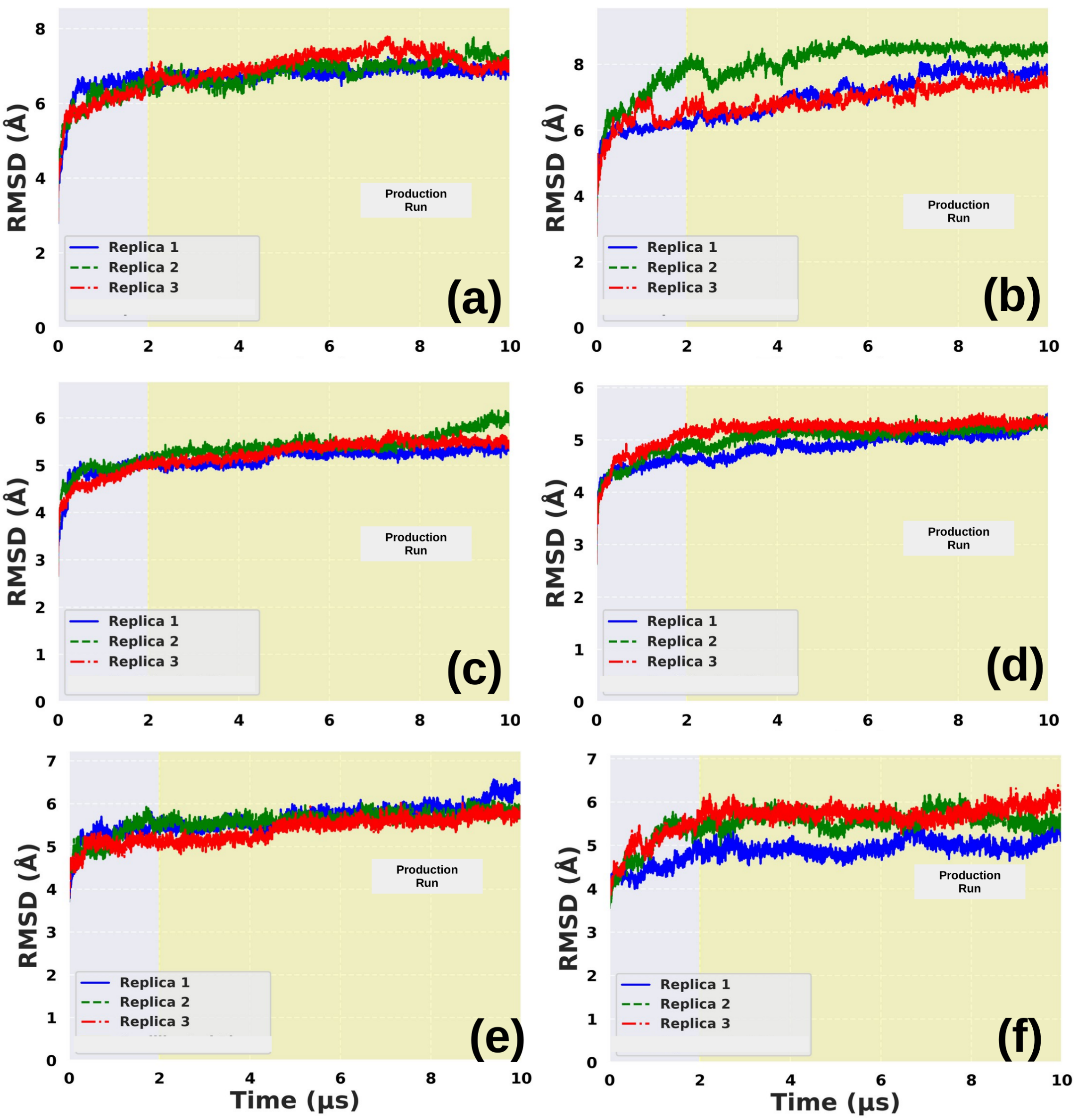


Fig. S3: RMSD plots for different CG-MD replicas: (a, b) histone core with the histone tails in the CENP-A nucleosome and CENP-A + CENP-N nucleosome systems, respectively; (c, d) histone core without the tails considered; (e, f) DNA in the CENP-A-containing nucleosome and in the CENP-A + CENP-N nucleosome systems. Replicas are shown in different colors. The last 8 $\mu$s of the trajectory were analyzed and depicted as production run, excluding the initial 2 $\mu$s for equilibration.

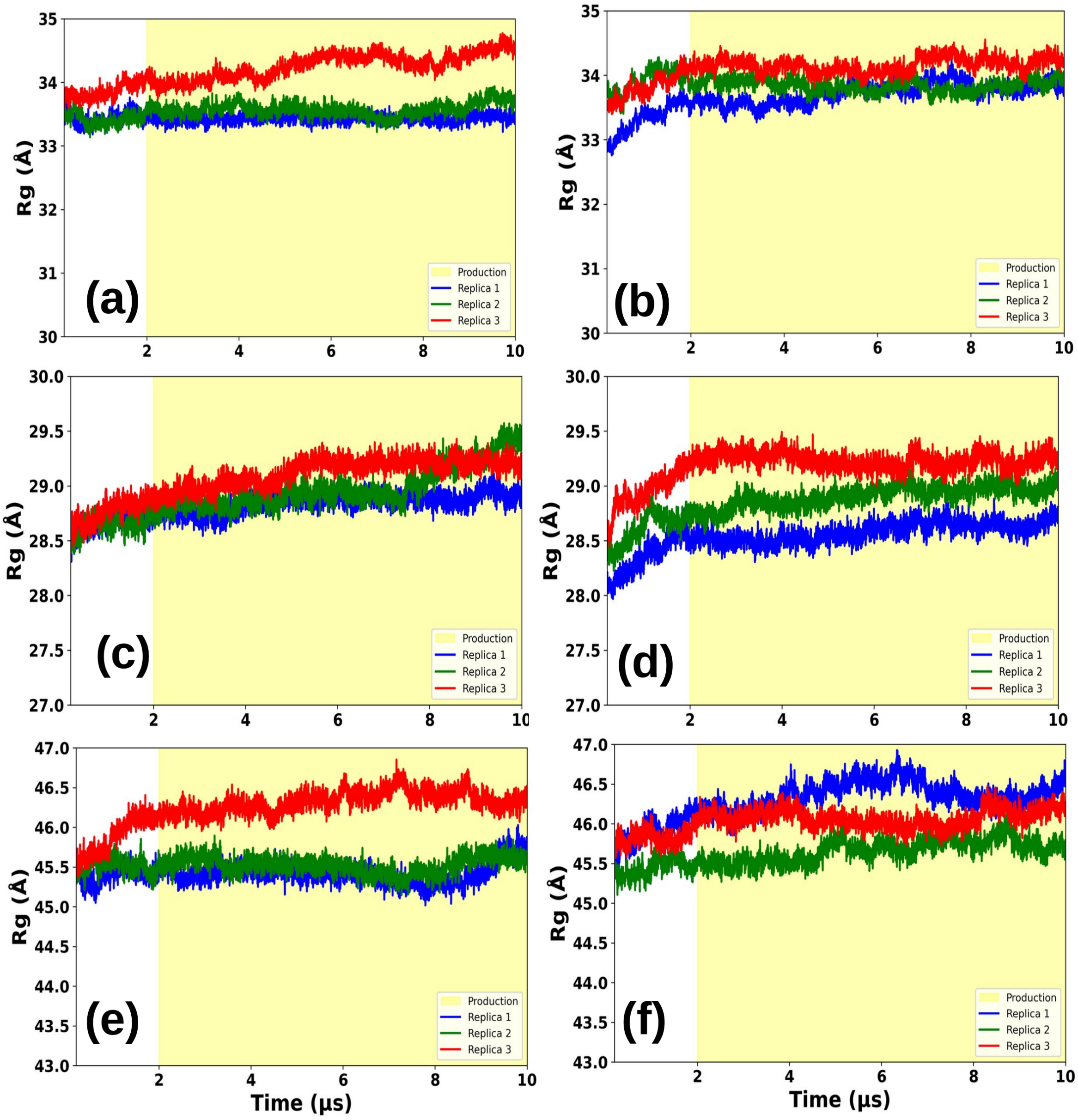


Fig. S4: Rg plots for different CG-MD simulations: (a, b) histone core with the histone tails in the CENP-A nucleosome and CENP-A + CENP-N nucleosome systems, respectively; (c, d) histone core without the tails considered; (e, f) DNA in the CENP-A-containing nucleosome and in the CENP-A + CENP-N nucleosome systems. Replicas are shown in different colors. The last 8 $\mu$s of the trajectory were analyzed and depicted as production run, excluding the initial 2 $\mu$s for equilibration.

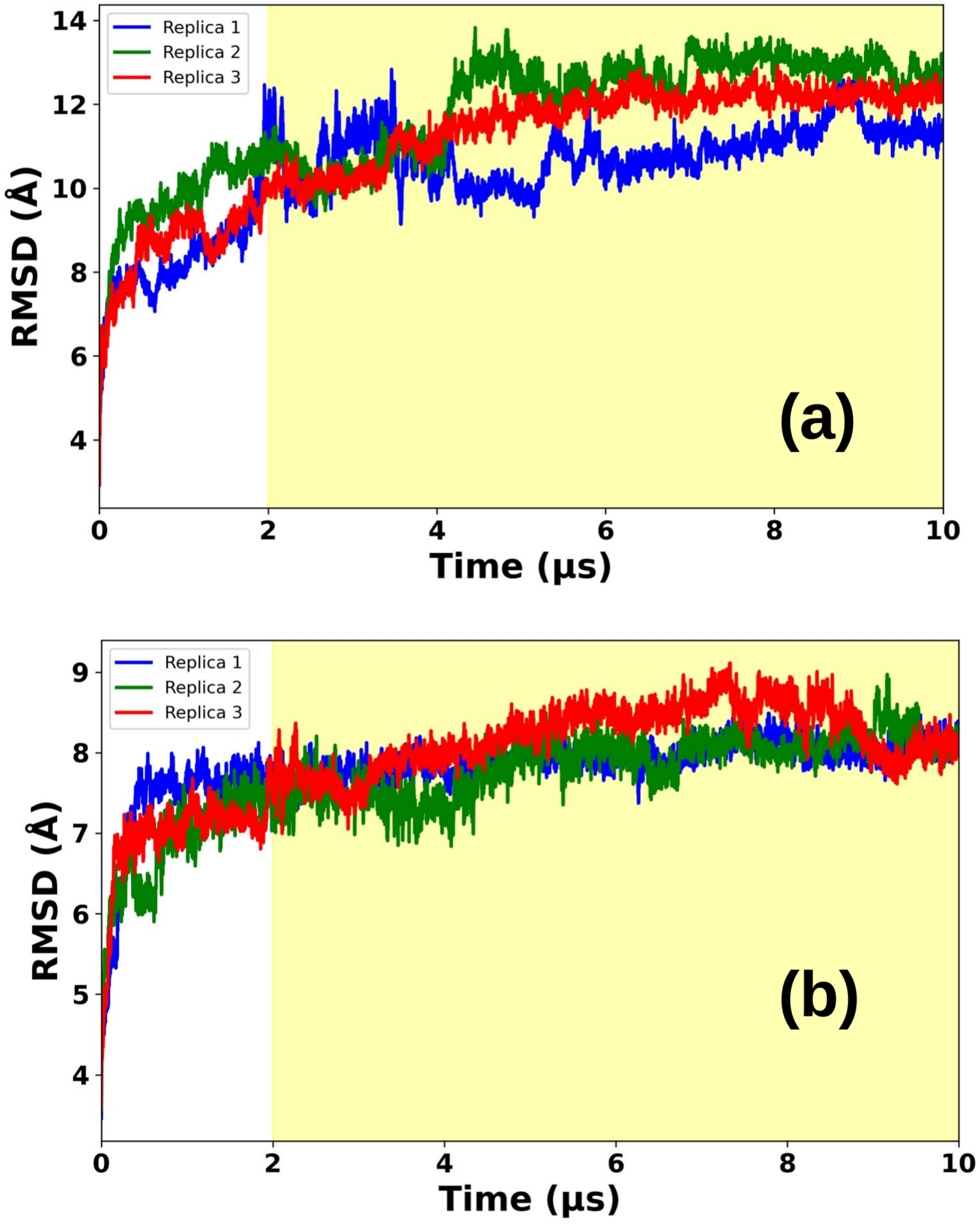


Fig. S5: RMSD plots calculated for histone tails in the (a) the CENP-A nucleosome and (b) CENP-A + CENP-N nucleosome systems, respectively. Replicas are shown in different colors. The last 8 $\mu$s of the trajectory were depicted as production run, excluding the initial 2 $\mu$s for equilibration.

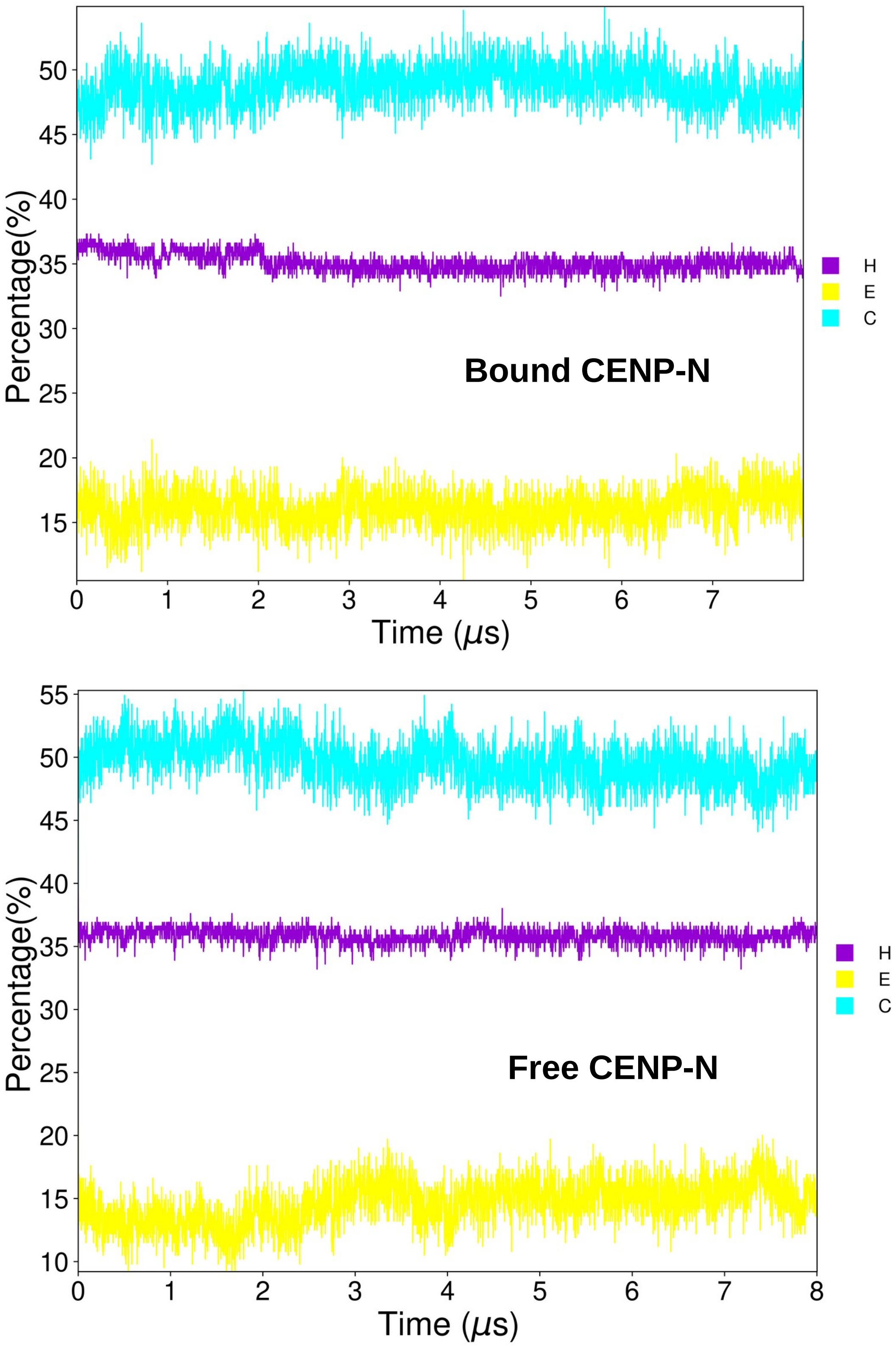


Fig. S6: Time evolution of the average secondary structure (in %) for both bound and free states of the CENP-N protein over the last 8 $\mu$s of the trajectory.

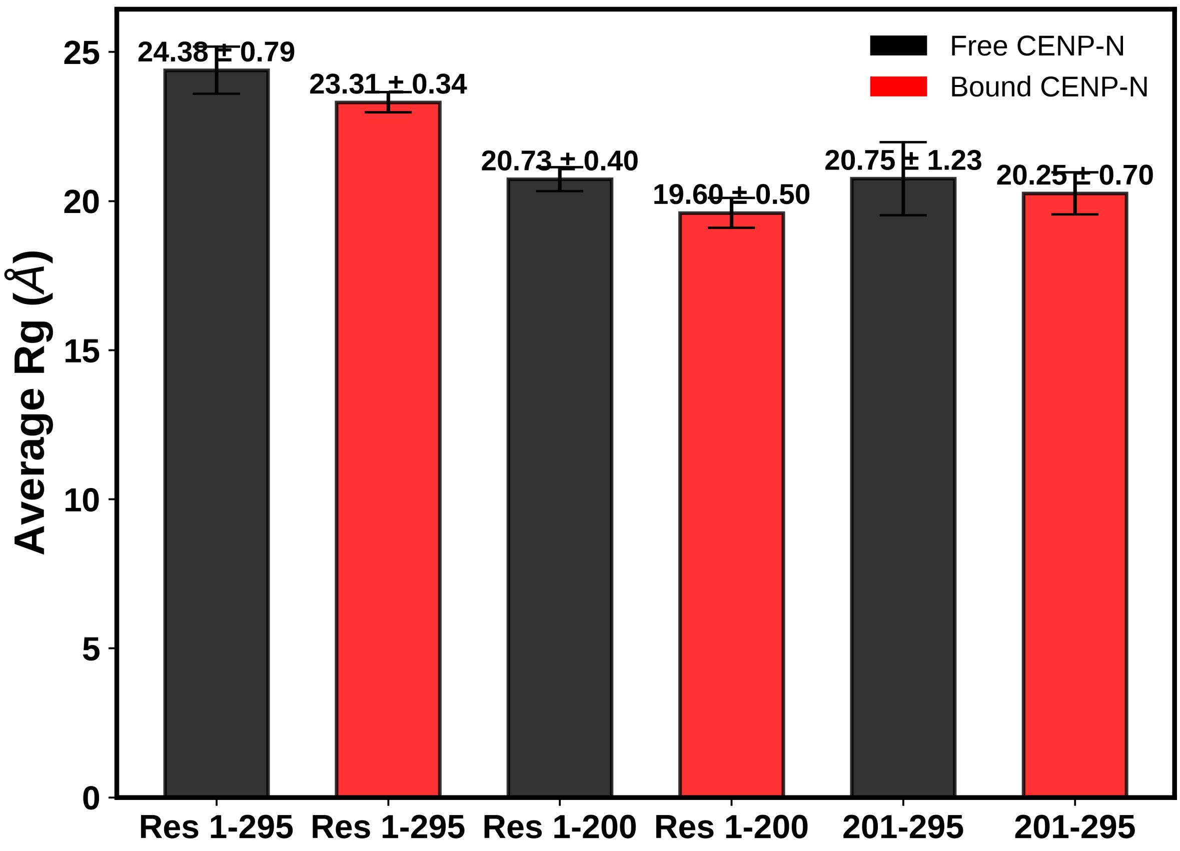


Fig. S7: The averaged Rg plots of the free and bound states of CENP-N calculated for three sets of residue regions: full sequence (Res 1-295), residues 1–200, and residues 201–295. Error bars represent standard deviations from three independent replicas.

.

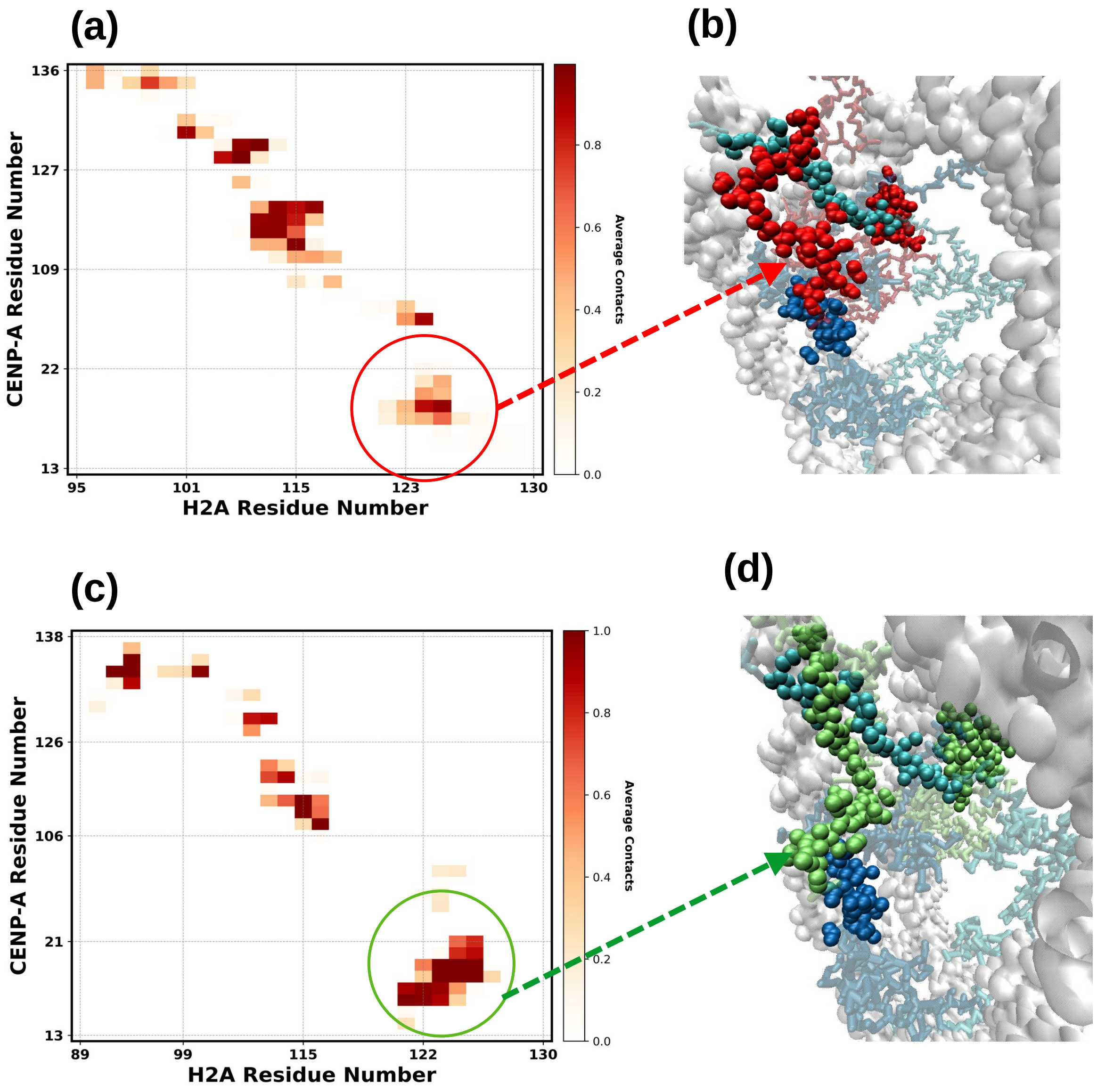


Fig. S8: (a) Contact map showing interactions between CENP-A and H2A residues in the presence of CENP-N. (b) Simulated structure highlighting CENP-A regions: residues forming contacts with H2A are shown in red, non-contacting regions are in cyan, and N-terminal residues with no contacts are in blue. (c) Contact map showing interactions between CENP-A and H2A residues in the absence of CENP-N. (d) Structural representation of CENP-A where residues in contact with H2A are marked in green. Color coding of other regions follows panel (b). The N-terminal region exhibiting significant changes in average contacts with H2A is circled for emphasis.

.
